# Adaptive immunity is dispensable for appendage regeneration in highly regenerative vertebrates

**DOI:** 10.64898/2026.04.13.718128

**Authors:** Lizbeth Airais Bolaños Castro, Yaswanth Othayoth Valappil, Andreas Petzold, Franziska Knopf, Maximina H. Yun

**Affiliations:** Technische Universität Dresden, Center for Regenerative Therapies TU Dresden (CRTD); Dresden, Germany; Dresden-concept Genome Center, DFG NGS Competence Center, c/o Center for Molecular Bioengineering (CMCB), TU Dresden; Dresden, Germany; Center for Healthy Aging, Faculty of Medicine Carl Gustav Carus, Dresden, Germany; Physics of Life Excellence Cluster TU Dresden; Dresden, Germany; Chinese Institutes for Medical Research, Beijing, China

## Abstract

Adaptive immunity has been implicated in tissue repair and homeostasis, however its requirement for complex appendage regeneration in adult vertebrates remains unknown. Here, we show that adaptive immune components are dynamically recruited to regenerating appendages. Using genetic lymphocyte ablation in highly regenerative vertebrates, axolotl (*Ambystoma mexicanum*) and zebrafish (*Danio rerio*), we show that mature T and B cells are dispensable for limb, tail and fin regeneration in sexually mature animals. Despite depletion of peripheral and lymphoid T and B populations, Rag1−/− axolotls and zebrafish regenerate appendages with normal kinetics, patterning, and skeletal outcomes. Rag1 −/− regenerating blastemas undergo transcriptomic remodelling including alterations in innate immune and extracellular matrix remodelling genes, accompanied by enhanced neutrophil/myeloid infiltration, highlighting innate immunity as a potential compensatory element for regenerative success. Together, these results indicate that adaptive immunity is not required for restoration of complex appendages in vertebrates, a finding of basic and translational relevance.

**Highlights:**

- Rag1−/− axolotls lack mature T and B cells in lymphoid organs and periphery.
- Rag1−/− axolotls and zebrafish regenerate appendages with kinetics, patterning and sizes comparable to wild-type siblings.
- Rag1−/− blastemas show downregulation of adaptive immune programs, modulation of innate immune genes, and heightened myeloid activity and/or infiltration in Rag1−/− animals.
- Innate immune compensation likely enables functional regeneration in the absence of mature adaptive lymphocytes.

## Introduction

Vertebrate appendage regeneration entails a coordinated program of wound healing, cellular dedifferentiation, progenitor proliferation and patterned redifferentiation that restores complex multi-tissue structures (Tanaka 2011; Cox 2019; Sehring & Weidinger 2022). In highly regenerative vertebrate species—urodele amphibians and teleost fish—this process proceeds through a blastema of proliferating progenitors that arises beneath a specialized wound epidermis (Tanaka 2011). Successful regeneration requires not only intrinsic plasticity of parenchymal and stromal cells but also the timely engagement of immune processes that clear debris, restrain infection, remodel extracellular matrix (ECM) and provide trophic cues that shape progenitor behavior (Bolaños-Castro 2021; Cox 2019).

Historically, an inverse relationship between adaptive immune sophistication and regenerative capacity has been proposed: many invertebrates that lack canonical adaptive immunity retain whole-body regenerative abilities, fish and salamanders with comparatively rudimentary adaptive arms regenerate extensively, and adult mammals—whose adaptive responses are highly developed—exhibit limited regenerative potential (Frippiat 2001). Yet this binary view has been supplanted by a more nuanced picture in which immunity is not merely permissive or inhibitory, but a context-dependent regulator of regeneration (Bolaños-Castro 2021; Cox 2019). Accumulating evidence indicates that immune components are essential for regeneration across taxa, with both innate and adaptive components being central to complex regenerative processes (Bolaños-Castro 2021; Godwin 2013; Yun 2015; Wynn & Vannella 2016).

Innate immunity provides indispensable early signals after injury (Bolaños-Castro 2021). Complement activation, proinflammatory cytokines and chemokines rapidly recruit neutrophils and monocyte-derived macrophages (Bolaños-Castro 2021; Godwin 2013). These populations execute debris clearance, secrete matrix metalloproteinases (MMPs) and angiogenic factors, and orchestrate transitions from pro- to anti-inflammatory states that permit blastema formation (Godwin 2013; Debuque 2021). Functional studies across vertebrates indicate a requirement for innate immunity in complex regeneration, including zebrafish tail fin (Petrie et al, 2014; Morales Allende 2019), fin fold (Hasegawa et al, 2017; Nguyen-Chi et al, 2015; Miskolci et al, 2019; Nguyen-Chi et al, 2017), bone (Geurtzen et al, 2022) and brain (Kyritsis et al, 2012), lizard tail (Londono et al, 2020), frog (Aztekin et al, 2020), and mouse digit tip and ear pinna (Simkin et al, 2017a and b). Single-cell profiling has revealed temporally distinct myeloid populations in axolotl (*Ambystoma mexicanum*) blastemas, including recruited Trem2+ macrophages and liver-derived macrophage contributions (Leigh et al, 2018, Rodgers et al, 2020). Clodronate-mediated macrophage depletion in salamander species impairs limb (Godwin 2013), heart (Godwin et al 2017) and lens (Tsissios et al 2024) regeneration through disruption of ECM remodelling, reduction of proliferation and dysregulation of signalling pathways. Further, macrophages orchestrate phagocytic clearance of senescent cells during axolotl limb regrowth, a process necessary for normal progression (Yun et al, 2015). Neutrophils appear early and transiently prior to blastema formation, consistent with a role in initial wound conditioning, but their absolute requirement in salamander regeneration remains unresolved (Rodgers et al, 2020).

Adaptive immunity contributes additional, often critical tissue-specific layers of regulation. Across vertebrates, subsets of T lymphocytes exert pro-regenerative functions that extend beyond canonical antigen-specific roles (Charlemagne 1979). In mice, combined T and B cell depletion impairs redifferentiation after pancreatitis (Folias 2014), and T cells modulate collagen deposition and functional recovery after fracture (El Khassawna 2017). B lymphocytes can enhance opsonization, modulate neutrophil and macrophage activity via antibodies and cytokines, and contribute to angiogenesis and remodelling in some contexts (Folias 2014; El Khassawna 2017). Further, regulatory T cells (Tregs) are critical for murine lung injury resolution (D’Alessio et al, 2009), wound healing (Nosbaum et al, 2016) and hair follicle growth (Ali et al, 2017). In zebrafish, FOXP3+ Tregs have been shown to secrete growth factors -Igf1, Nrg1, Ntf3, amphiregulin- that directly stimulate proliferation of resident progenitors and promote anti-inflammatory environments conducive to repair in a tissue-specific manner, modulating spinal cord, heart, brain (Hui et al, 2017) and caudal fin regeneration (Hui et al, 2022).

Despite identification of T- and B-cell markers in regenerating structures, functional dissection of adaptive lymphocytes in salamander regeneration is still limited (Frippiat et al, 2001). Nevertheless, pharmacological immunosuppression experiments (Fahmy et al, 2002) and recent single-cell analysis led to the prevalent view that adaptive immune cells contribute to complex appendage regeneration (Leigh et al 2018; Rodgers et al 2020).

Here, we directly probe the role of adaptive lymphocytes in appendage regeneration. We profile lymphopoietic gene expression in the axolotl, track T and B cell dynamics in regenerating appendages, and leverage genetic models of lymphocyte deficiency via targeted disruption of Rag1 and FoxN1 to determine how absence of mature adaptive cells affects limb and tail regrowth. We further address whether Rag1 is required for zebrafish caudal fin regeneration. We uncover that adaptive immunity is dispensable for successful appendage regeneration, challenging prior notions and refining our understanding of immune–regenerative interactions, with implications for both basic biology and translational approaches.

## Results

### T and B cell ontogeny in the axolotl

While important molecular features of the axolotl innate immune system have been described (Godwin, 2013; Lopez et al, 2014), those corresponding to the adaptive branch have only recently begun to be elucidated. To characterise the axolotl adaptive immune system, we leveraged our recently published single cell Atlas of the axolotl thymus (Czarkwiani, Lobo, Bolaños Castro et al, 2025) to generate a panel of *in-situ* hybridization (ISH) probes for key adaptive cell markers (Supplementary Table 1) enabling us to assess the ontogeny of adaptive immunity in this species. The analysis focused on post-hatching expression of key markers in the three main hematopoietic organs in axolotls (Lopez et al, 2014, Czarkwiani, Lobo, Bolaños Castro et al 2025), namely liver, spleen and thymus. Expression of hematopoietic markers *Flt3* and *Runx1* is present in the cortical area of the liver at 3 weeks of age (Supplementary fig. 1), while only observed in spleen after 1 month, supporting previous findings indicating the liver is the first organ with hematopoietic potential (Lopez et al, 2014). By 1 month of age, both liver and spleen show expression of lymphocyte markers *Rag1* and *Cd79b* (B cell receptor complex-associated protein β chain), while spleen also exhibits expression of the class M B cell marker *Igmhc*, and *Flt3, Runx1, Rag1* and *Trac* are expressed in the thymus (Supplementary fig. 1). These results indicate that, upon hatching −3 weeks post-fertilization-, axolotls already exhibit signs of haematopoiesis, while expression of lymphopoietic genes ensues at 1 month of age.

Next, we asked whether hematopoietic organs sustain lymphopoiesis after sexual maturation. Livers of 1-year old axolotls show expression of *Rag1*, *Cd79b, Cd4* and *Cd8b,* whereas spleens express *Rag1*, *Lck, Cd79b, Cd4, Cd8b* and *Igmhc,* and thymic nodules retain expression of *Flt3, Runx1, Rag1* and *Trac* (Supplementary fig. 2), indicating that all three organs retain the ability to support lymphopoiesis.

### T and B cells are dynamically localised to regenerating axolotl limbs

While T and B lymphocytes have been identified within regenerating axolotl limbs (Leigh et al, 2018; Rodgers et al, 2020), a detailed, quantitative assessment of adaptive cell recruitment dynamics during appendage regeneration is lacking. To this end, we performed anti-CD2 (T cells) or anti-CD79A (B cells) immunofluorescence staining and tissue clearing coupled with light-sheet microscopy imaging at different regeneration stages in 5-cm-long animals (SVL) (Figure 1). This approach allows for the imaging of entire tissue structures at the millimetre scale such as the axolotl limb, enabling quantitation at single cell resolution (Subiran Adrados et al., 2021). Both T and B cells are present in the regenerating limb, albeit with differences in spatial distribution (Figure 1a). Both T and B cells are localised along the skin and at the distal-most region of the regenerate, in contrast to the more homogeneous distribution of both populations in homeostasis (Figure 1a). Quantification of the total cell number per volume (cell density) within the regenerating tissue unveiled two peaks of CD2+ T cell accumulation, firstly at the onset of wound healing (1dpa and 2dpa time points) and the early blastema formation, and secondly at the digit outgrowth stage (Figure 1b). In contrast, CD79A+ B cell density peaks during the wound healing process and afterwards remains mostly constant along regeneration (Figure 1b). Quantification of thymidine nucleotide analogue EdU incorporation after a two-hour pulse combined with CD2 or CD79A immunofluorescence revealed no significant increases in cell proliferation (data not shown) suggesting that the increase in T and B cells during regeneration is due to cell homing. The presence of T and B lymphocytes at critical stages of appendage regeneration raised the possibility of their functional involvement in this process.

**Figure 1.**
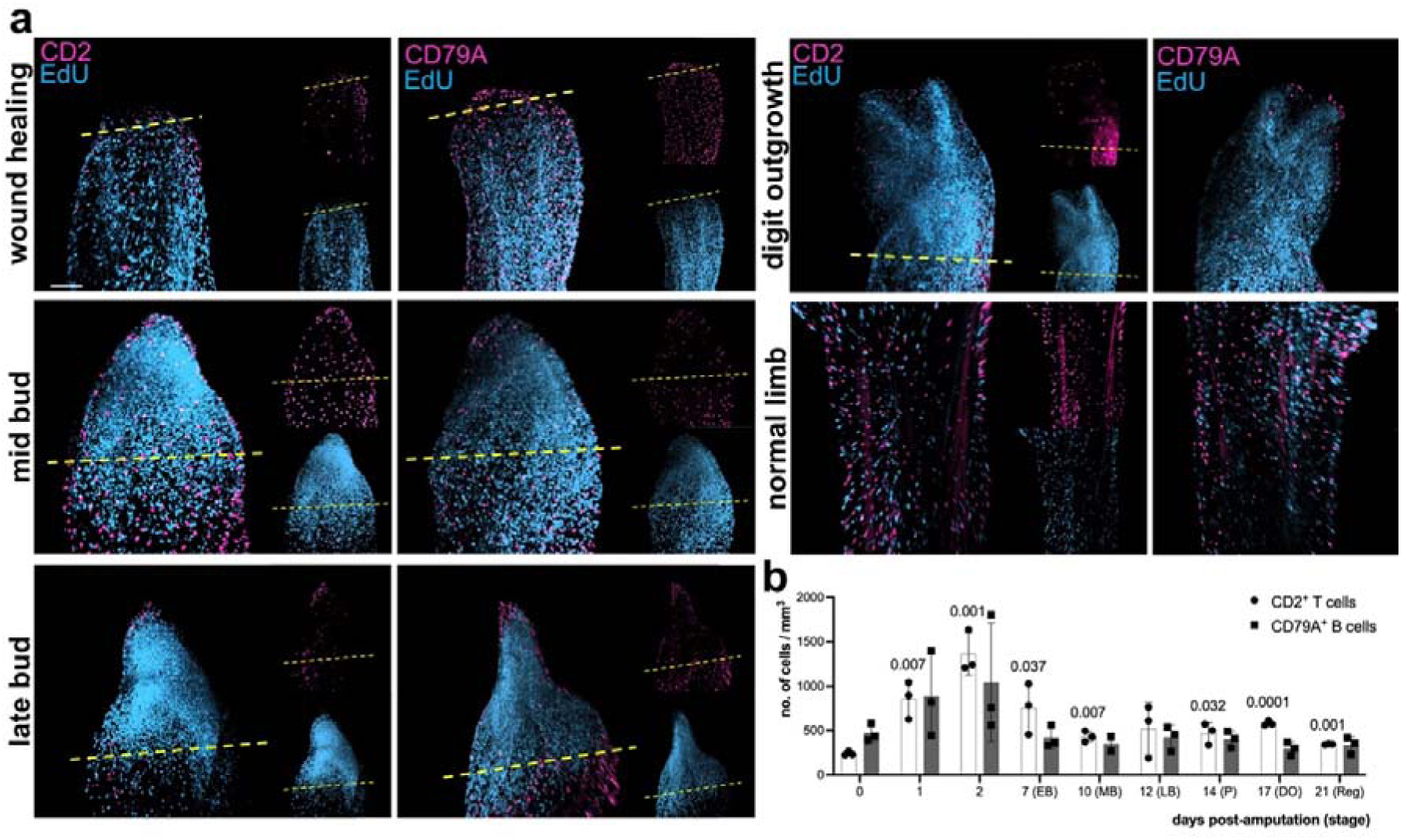
CD2+ T- and CD79A+ B-lymphocytes locate at the regenerating limb. a) 3D reconstructions of whole-mount immunofluorescence coupled with EdU stainings and tissue clearing across regeneration for CD2 (left panels) or CD79A (right panels). Scale bar = 200 μm. b) CD2+ or CD79A+ cells per volume within the regenerating tissue. n=3, mean with SD, unpaired Student’s T-test of each time point against untouched (0) control. Bars without p-values showed no statistical significance against untouched control.

### Axolotl *Rag1*−/− exhibit loss of mature T and B cells

To genetically probe the requirement for T and B cells during axolotl appendage regeneration, we generated a mature lymphocyte depletion model via disruption of the RAG1 recombinase (Mombaerts et al, 1992). Targeting the axolotl *Rag1* locus with CRISPR/Cas9 using gRNAs directed to exon 2 generated a series of indel alleles that produced frameshifts and predicted early stop codons (Supplementary fig. 3a, b). Sequencing of mutant alleles from founder animals revealed deletions of 1–11 nucleotides that truncate the RAG1 open reading frame at multiple upstream positions. Adult Rag1−/− animals are viable, fertile and reach sexual maturity without gross developmental abnormalities, allowing assessment of regeneration in sexually mature contexts. Of note, F1 and F2 animals were used for all subsequent *Rag1*−/− line assessment.

Whole-mount nuclear staining and 3D reconstructions showed significantly reduced thymic nodule volume in *Rag1* knockouts compared with heterozygous siblings (Fig. 2a), with reductions being more modest yet in line with observations in mice (Mombaerts et al, 1992). Spleens from *Rag1*−/− animals showed reduced mass and altered histoarchitecture, with spleen weight significantly lower in both sexes (Fig. 2b). Gene expression profiling of spleen tissue revealed marked downregulation of B cell markers *Cd79b* and *Igmhc* (non-detectable in most knockout samples) as well as *Rag1* itself, while thymic tissue exhibited strong reductions in T cell markers *Trac*, *Cd4* and *Cd8b* (Fig. 2c–d). Blood samples mirrored these deficits with decreased *Rag1*, *Trac* and *Igmhc* expression (Fig. 2f). Importantly, the thymus of *Rag1*−/− animals exhibits a near absence of CD3+ T cells (Fig. 2e), indicating the successful depletion of mature T lymphocytes in the knock-out line. Further, immunostaining corroborated that CD79A+ B cells and CD3+ T cells are absent or dramatically reduced in *Rag1*−/− limbs (Fig. 2g, 2h). Collectively, these data demonstrate efficient elimination of mature adaptive lymphocytes and thymic hypoplasia in the axolotl *Rag1*−/− that parallels *Rag1* deficiency in other vertebrates, and provide a suitable model to explore regeneration in the absence of adaptive immunity.

**Figure 2.**
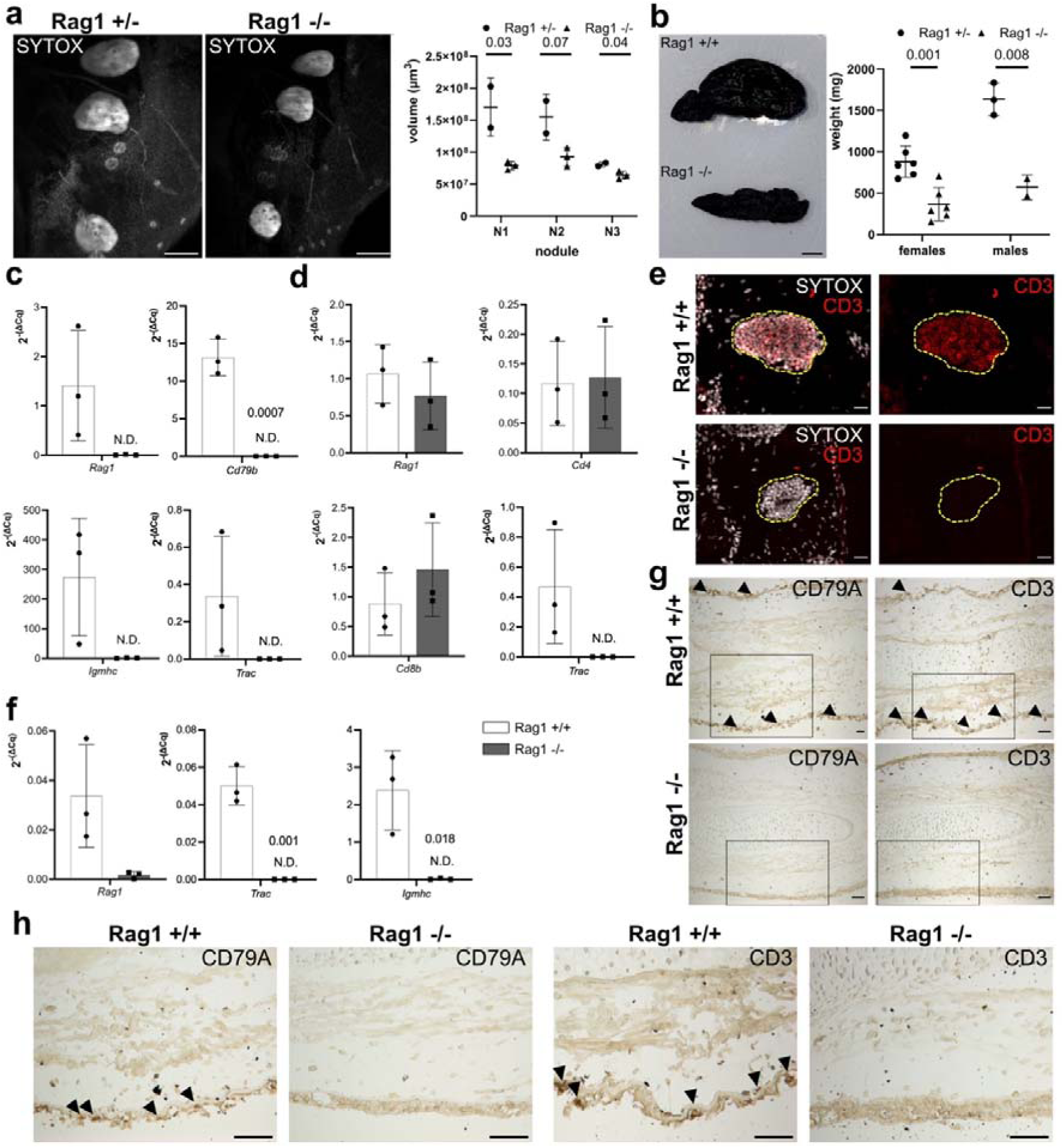
Rag1 knockout depletes mature T and B lymphocytes in hematopoietic tissues and periphery. a) Left: maximum projection of whole-mount nuclear staining of optically cleared heads from Rag1 −/− and heterozygous sibling animals. Scale bar = 500 μm. Right: thymic nodule volume (n=2 for HT, n=3 for KO). b) Left: spleen from Rag1 −/− and heterozygous sibling animals. Scale bar = 500 μm. Right: spleen weight separated by sex (females: n=6, males: n=3 for HT, n=2 for KO). c) *Rag1*, *Cd79b*, *Igmhc* and *Trac* gene expression in splenic tissue (n=3). d) *Rag1*, *Cd4*, *Cd8b* and *Trac* gene expression in thymic tissue (n=3). e) CD3 immunofluorescence in thymic tissue. Scale bar = 50 μm. f) *Rag1*, *Igmhc* and *Trac* gene expression in blood (n=3). g) Immunohistochemistry against CD79A or CD3 on limb cryosections. Scale bar = 100 μm. h) Higher magnification of insets in (g). Scale bar = 50 μm. Positive cells indicated with black arrowheads. n=3 for (a-h). ND: not detected. Unpaired Student’s T-test performed for all statistical analyses against the wildtype or heterozygous data. Bars without p-values showed no statistical significance against controls.

### Regenerative capacity is preserved in *Rag1*−/− axolotls and zebrafish

To directly test whether mature lymphocytes are required for complex appendage regeneration, we performed limb amputations (Figure 3a) or tail fin clips (Figure 3b) on sexually mature *Rag1*−/− and (heterozygous and) wild-type siblings (axolotls 2-year-old). Macroscopic imaging and quantitative area measurements showed no significant differences between genetic backgrounds in timing or extent of limb and fin regrowth over the measured interval (Fig. 3a-d). Regenerates in *Rag1*−/− axolotls progressed through canonical regenerative stages (wound healing, early bud, mid-bud, palette/digit formation) with similar proportions of animals at each stage at corresponding time points (Fig. 3c,d). To test whether this outcome generalises across taxa, we analysed tail fin regeneration in 8-month-old *Rag1*−/− zebrafish and observed comparable regenerative rates and lengths relative to wild-type and heterozygous siblings (Fig. 3e), further indicating that mature T and B cells are dispensable for regeneration. Juvenile axolotls lacking *Rag1* also regenerate limbs and tails without delay (Supplementary fig. 4). Finally, a recently established *Foxn1*−/− axolotl line which exhibits T cell deficiency (Czarkwiani, Lobo, Bolaños Castro et al, 2025), exhibits no apparent defects in neither limb nor tail regeneration (Supplementary fig. 5). These cross-appendage, cross-species results indicate that mature adaptive lymphocytes are not required for the initiation or progression of complex appendage regeneration in highly regenerative vertebrates.

**Figure 3.**
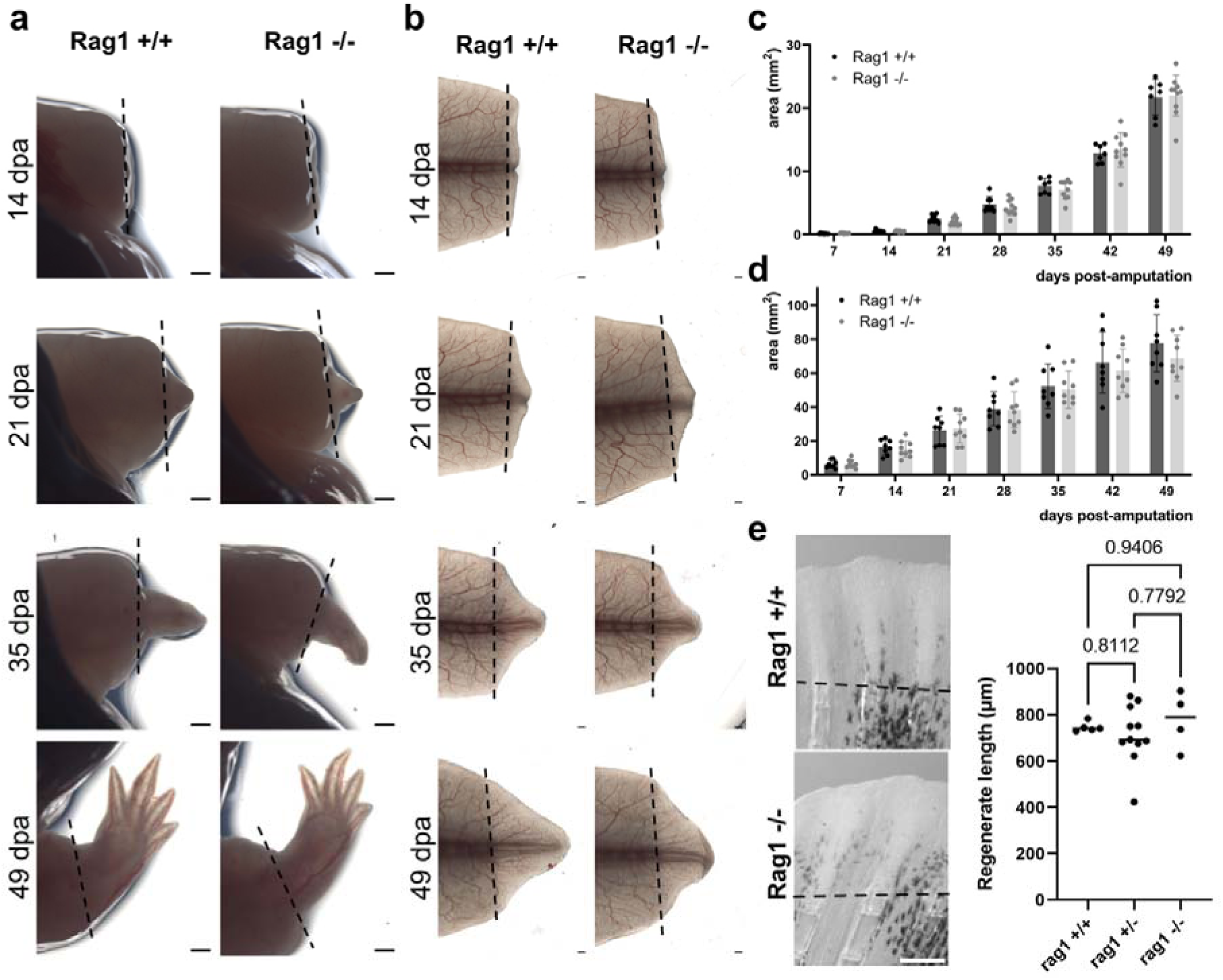
Limb and tail regeneration is retained in sexually mature Rag1−/− axolotls and zebrafish. a,b) Limb (a) and tail (b) regeneration in 2-years-old wildtype and Rag1 −/− sibling axolotls. Scale bar = 1 mm. c,d) Area of regenerated limb (c) or tail (d) tissue (n=8 for WT, n=10 for KO, N=2, mean with SD). e) Left: tail fin regeneration in 8-months-old wildtype and Rag1 −/− zebrafish. Scale bar = 250 μm. Right: length of regenerated tail fin at 4 dpa (n=5 for WT, n=11 for HT, n=4 for KO). Unpaired Student’s T-test performed for statistical analyses against the wildtype data in (c,d). No statistical significance observed. Brown Forsythe and Welch ANOVA test for data in (e).

### Normal morphogenesis and skeletal regeneration in the absence of adaptive lymphocytes

Although gross regrowth was intact, adaptive immunity could potentially be required for accurate patterning or formation of skeletal elements. We therefore examined regenerated skeletal structures by alcian blue staining of cartilage in axolotl limbs and by immunostaining for Alcama (Activated leukocyte adhesion molecule a) labelling osteoblasts in regenerating zebrafish fins. *Rag1*−/− axolotls reconstructed humerus, radius, ulna and digit cartilages that were morphologically indistinguishable from wild-type regenerated structures (Fig. 4a).

**Figure 4.**
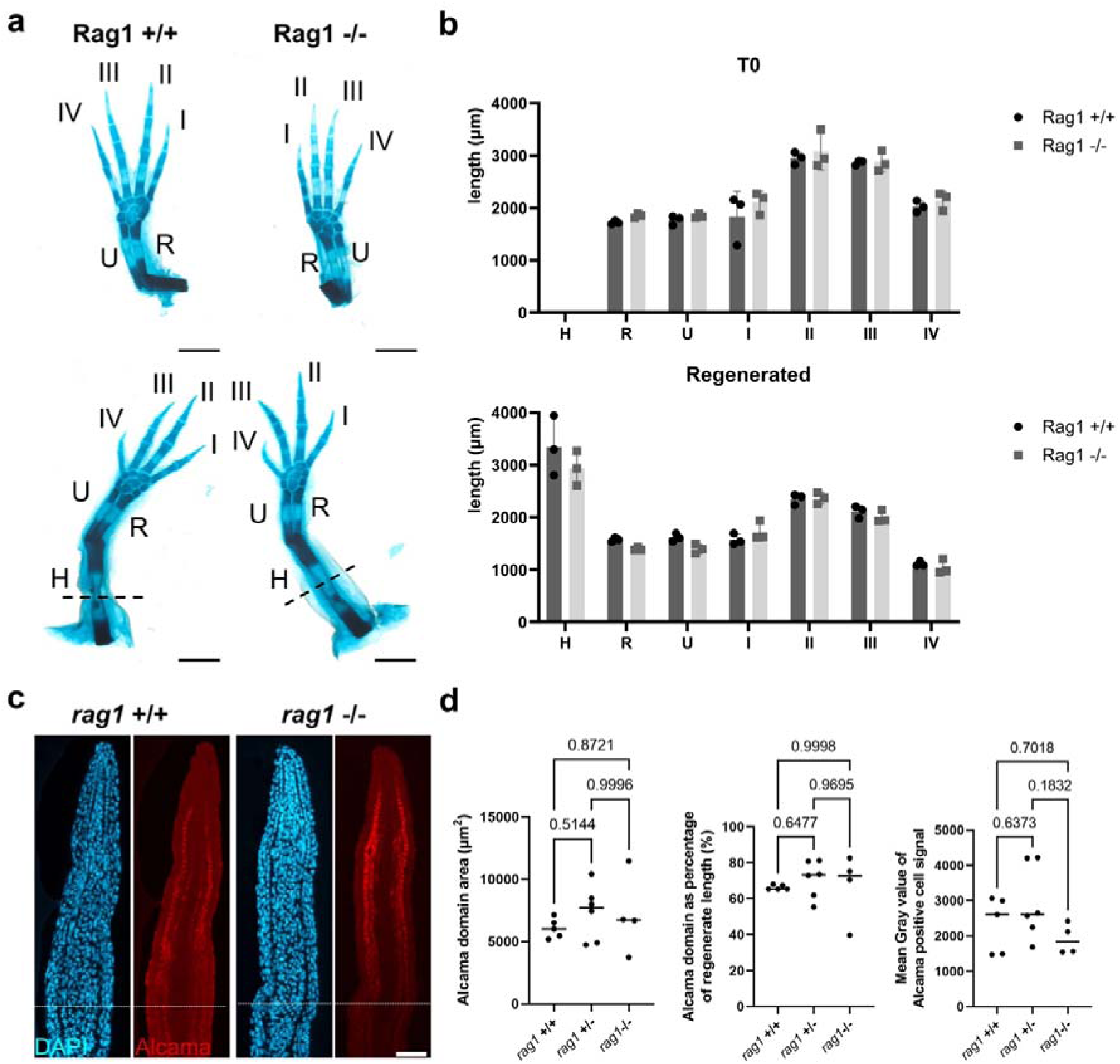
Lymphocyte-depleted tissues regenerate skeletal structures without defects. a) Alcian blue skeletal stainings on uninjured (top) and regenerated (bottom) limbs from wildtype and Rag1 −/− sibling axolotls. Skeletal elements are indicated with uppercase letters. H: humerus, R: radius, U: ulna, I-IV: digits 1 to 4 (n=3). Scale bar = 1 mm. b) Length of developed and regenerated skeletal structures (n=3). Unpaired Student’s T-test performed for statistical analyses against the wildtype data. No statistical significance observed. c) IF for Alcama on regenerating zebrafish tail fins from wildtype and Rag1 −/− siblings at 4 dpa. Scale bar = 50 µm. d) Alcama positive area, percentage of Alcama domain and intensity of Alcama signal (n=5 for WT, n=6 for HT, n=4 for KO). Brown Forsythe and Welch ANOVA test.

Quantitative measurements of individual skeletal element lengths in both intact (T0) and regenerated limbs revealed no significant genotype differences (Fig. 4b), indicating conserved scale and proportion. In zebrafish fins, Alcama domain area and signal intensity in regenerating blastemas were comparable across genotypes (Fig. 4c-d). Thus, despite the absence of mature T and B cells, the morphogenetic programs that direct skeletal reformation and positional identity remained functional.

### Myeloid alterations and transcriptional remodeling in lymphocyte-depleted blastemas

To probe molecular consequences of adaptive immune loss during regeneration, we performed bulk RNA-seq on axolotl limb blastemas harvested at 12 dpa, a time point capturing active blastema proliferation and patterning (Fig. 5a). Differential expression analysis between *Rag1*−/− and control blastemas revealed two principal trends. First, several canonical adaptive immune and lymphocyte-associated genes were significantly downregulated in *Rag1*−/− blastemas (Fig. 5b-c; Supplementary fig. 6a-c). Gene Ontology (GO) enrichment on downregulated DEGs highlighted adaptive immune system processes, antigen receptor-mediated signalling and lymphocyte activation (Fig. 5c). Concurrently, immunostaining confirmed near absence of CD79A+ B cells and CD3+ T cells in *Rag1*−/− blastemas, mirroring lymphoid gene repression observed by RNA-seq (Fig. 5d).

**Figure 5.**
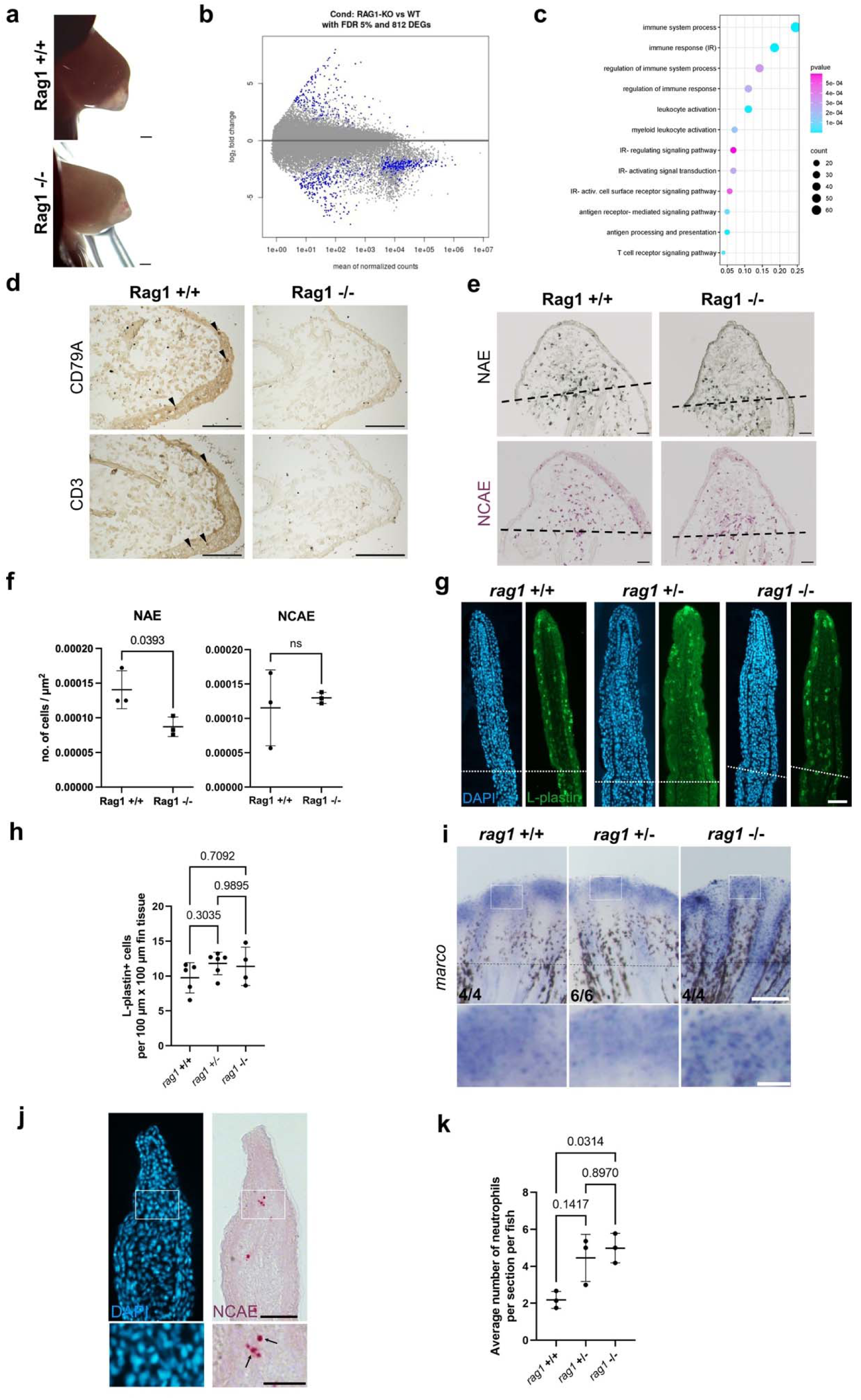
Lymphocyte-depleted blastemas exhibit differential infiltration of myeloid populations. a) Limb blastemas collected for the bulk-RNAseq dataset at 12dpa (n=3). Scale bar = 1 mm. b) Volcano plot of DEG analysis highlighting in blue the significantly up- and down-regulated genes. c) GO enrichment analysis on downregulated DEGs. d) IHC on limb blastema cryosections for CD79A or CD3 from wildtype and Rag1 −/− siblings (n=3). Scale bar = 100 µm. e) NAE or NCAE stainings on limb blastema cryosections. f) NAE or NCAE positive cells per area (n=3 sections per animal, n=3 animals). g) L-plastin IF on regenerating tail fins from wildtype, heterozygous and Rag1 −/− sibling zebrafish at 4 dpa. Scale bar = 50 µm. h) L-plastin positive cells per area (n=5 for WT, n=6 for HT, n=4 for KO). i) marco ISH on zebrafish tail fin blastema cryosections. Scale bar = 200 µm and 25 µm (insets). j) NCAE staining on zebrafish tail fin blastema cryosections. Scale bar = 50 µm and 25 µm (insets). k) Quantifications of NCAE+ cells normalized per section (n=3 fish per condition). Brown Forsythe and Welch ANOVA test for data in (h,k).

Second, *Rag1*−/− blastemas exhibited alterations of genes linked to innate immune responses. While the macrophage marker *Mpeg1* was downregulated in *Rag1*−/−, the neutrophil activity gene *Mpo* was significantly upregulated (Supplementary fig. 6d-e). GO enrichment reflected alterations in the regulation of reactive-oxygen species (ROS, Supplementary fig. 6d), which could result from increased neutrophil activity. Expression of key morphogenetic genes implicated in limb patterning and chondrogenesis was preserved or modestly upregulated in *Rag1*−/− samples, consistent with maintained patterning programs despite immune remodelling.

### Increased myeloid infiltration in lymphocyte-depleted blastemas

To validate transcriptomic indications of altered innate infiltration we performed histochemical and immunochemical assays on regeneration-stage cryosections. NAE/NCAE staining, markers of granulocytic/myeloid populations, revealed a decrease in macrophage (Figure 5e) and a mild yet non-significant increase in neutrophils in *Rag1*−/− axolotl limb blastemas (Figure 5e). Considering the aforementioned observations, it is likely that a qualitative rather than quantitative alteration in neutrophils is associated with RAG1 deficiency in axolotls. In line with this notion, anti L-plastin immunohistochemistry indicates no significant changes in leukocyte populations in *rag1*−/− zebrafish tail fin blastemas, however fins exhibit enhanced *marco* (*macrophage receptor with collagenous structure*, Jing et al, 2013) expression (Fig. 5 g-i) consistent with an increased innate immune response by macrophages and/or dendritic cells (Zakrzewska et al, 2010). Further, NCAE quantification showed statistically significant neutrophil increases in *rag1*−/− animals (Fig. 5j-k). The observed increases in myeloid activity and/or infiltration during appendage regeneration in *Rag1* mutants might reflect a compensatory response to adaptive deficiency and/or altered inflammatory signalling in the regenerating microenvironment.

## Discussion

Our cross-species *Rag1* genetic ablation analysis, and that of Umeano *et al*. (N. Leigh, personal communication), demonstrates that mature adaptive lymphocytes are not required for the successful regeneration of complex appendages in regeneration-competent vertebrates. *Rag1*−/− axolotls and zebrafish regenerate limbs, tails and fins, respectively, with preserved kinetics, patterning and skeletal restoration despite loss of mature T and B lineages. The principal system-level consequence of adaptive loss is a remodelling of the regenerative immune milieu toward heightened granulocytic/myeloid cell processes and/or infiltration and enhanced innate and ECM-associated transcriptional programmes, while the core morphogenetic programmes remain competent to restore structure and pattern.

The dispensability of mature T and B cells is unexpected in light of previous observations pointing towards the involvement of adaptive elements in both axolotls and zebrafish (Hui et al, 2017, Hui et al, 2022). Nevertheless, it is possible that such elements could contribute to subtler aspects not captured by our assays. These include the provision of antigen-specific regulation that limits chronic inflammation or autoimmunity after repeated injury, the mediation of immune memory that could regulate responses to subsequent insults, and effects on functional recovery (e.g., neuromuscular integration, fine-scale tendon/bone mechanical properties). Additionally, adaptive lymphocytes may influence regeneration under contexts of infection, metabolic stress or ageing, conditions not exhaustively modelled here and worthy of further consideration.

It is worth noting that innate-like lymphocytes (e.g., some innate lymphoid cells, gamma-delta-like T cells, or NK (natural killer)-like populations) may not depend on *Rag1* (Vivier et al, 2018) and could persist to varying degrees. These cells can produce cytokines and growth factors that overlap functionally with adaptive lymphocytes (Spits et al, 2016). Further, stromal and endothelial compartments can produce chemokines that recruit and polarize innate cells (Mueller & Germain, 2009). Thus, residual non–*Rag1*-dependent lymphoid or stromal-mediated signalling may contribute to the compensatory responses we observe.

Definitive attribution of regeneration-promoting functions to specific cell types will require single-cell resolution profiling and targeted depletion/lineage-ablation approaches.

Our data support a model in which regeneration-permissive tissues deploy innate immune mechanisms that can compensate for the absence of mature adaptive responses. Increased numbers of NCAE positive cells (zebrafish) and transcriptional enrichment for neutrophil (e.g., MPO, axolotl) and innate (e.g., *marco*, zebrafish) immune functions in *Rag1*−/− blastemas indicate augmented myeloid activity. This is consistent with reported increases in myeloid infiltration in retina, muscle and skin in *rag1−/−* adult fish (Kryshnavajhala et al, 2017; (Novoa et al, 2019). Myeloid cells are established mediators of debris clearance, cytokine, growth factor and protease secretion and matrix remodelling, processes critical to wound resolution and blastema formation (Godwin & Rosenthal, 2014). Enhanced or rebalanced myeloid responses in *Rag1*−/− animals likely provide the necessary microenvironmental cues to support dedifferentiation, proliferation and patterning of progenitor cells.

Concomitantly, expression of key developmental and ECM genes required for chondrogenesis and skeletal patterning was preserved or upregulated in *Rag1*−/− axolotl blastemas, indicating that tissue-intrinsic regenerative effectors remain intact and can operate within an altered inflammatory context. This highlights two complementary features of regenerative systems. First, the plasticity of the immune compartment, and particularly of the innate arm, to reconfigure in response to adaptive deficiency. Second, the inherent robustness of the developmental programs that drive patterning and redifferentiation, allowing them to proceed under modified immune conditions.

Our findings do not contradict the established importance of immune processes in regeneration, rather they refine the understanding of which immune components are strictly required in permissive species. In particular, they are consistent with previous reports demonstrating normal limb regeneration following splenectomy in axolotls (Fini & Sicard, 1980; Debuque et al, 2021) and transient reduction in fin regenerate length upon *foxp3*+ cell ablation in zebrafish (Hui et al., 2017). In mice, while B cells mediate cardiomyocyte proliferation during neonatal heart repair, B cell depletion actually improves recovery after injury (Tan et al, 2023). This suggests that lymphocytes are required in a context-dependent manner. It is likely that in an environment with toned-down adaptive responses, such as axolotl and zebrafish tissues, the requirements for mature lymphoid cells are limited.

Prior studies show that macrophage ablation impairs regeneration in salamanders (Godwin et al, 2013, Godwin et al, 2017) and zebrafish (Petrie et al, 2014), implicating innate populations as essential. The present work complements those results by showing that adaptive lymphocytes are dispensable in the contexts tested. Indeed, the apparent primacy of innate immunity for enabling blastema formation and progression is reinforced, whereas adaptive cells may serve modulatory roles (fine-tuning inflammation, antigen-specific regulation, long-term tissue homeostasis) that are not strictly necessary for the core morphogenetic program in these species.

Collectively, these results suggest that strategies aimed at harnessing or modulating innate immune responses -rather than adaptive suppression or augmentation per se- may be more directly relevant to promoting regenerative outcomes. Enhancing beneficial myeloid functions (pro-resolving macrophage phenotypes, timely neutrophil-mediated clearance, ECM-remodelling proteases in controlled contexts) or limiting maladaptive chronic inflammation might tip wound healing towards regeneration rather than fibrosis. However, the regenerative permissiveness observed here derives from species with intrinsic cellular plasticity; whether comparable innate-focused manipulations can overcome the more constrained plasticity of adult mammalian tissues remains an open question.

### Concluding remarks

Regeneration in permissive vertebrates is orchestrated predominantly by an innate-centric immune program that, together with robust tissue-intrinsic morphogenetic effectors, can accommodate the loss of conventional adaptive lymphocytes. While adaptive immunity can fine-tune outcomes and supply tissue-specific trophic factors in particular contexts, it is not an absolute requirement for the formation of a functional blastema or the reconstruction of appendages in axolotls and zebrafish. These conclusions sharpen our mechanistic understanding of immune-regenerative interactions and prioritise innate pathways as promising targets to unlock regenerative potential in less-permissive species.

### Limitations of the study

Several limitations temper interpretation. First, bulk transcriptomics averages signals across diverse cell types and cannot resolve cell-state–specific changes. Second, the morphological and skeletal assays employed herewith detect gross patterning defects but may miss microstructural, biomechanical or functional deficits. Third, *Rag1*−/− genetic backgrounds may harbour compensatory developmental adaptations; acute depletion models or temporally restricted ablation of adaptive cells could yield additional insights. Fourth, environmental variables (microbial exposure, husbandry) and injury contexts (infected vs sterile wounds, compounded injuries) can modulate immune-regenerative dynamics and were not exhaustively tested here. Finally, the contribution of residual innate-like lymphocyte subsets requires direct assessment.

## Materials and Methods Animal procedures

### Animal husbandry

All animal procedures and husbandry were conducted in accordance with the laws and regulations of the State of Saxony, Germany, and were approved by the ethics committee of Technische Universität Dresden and the Landesdirektion Dresden. Axolotls (*Ambystoma mexicanum*) used in this study were bred and maintained at the Axolotl Facility of the Center for Regenerative Therapies TU Dresden (CRTD), Germany.

Leucistic *d/d* strain axolotls, as well as mutant animals on a *d/d* genetic background were used for all experiments. Larvae were fed *Artemia* daily until they reached a snout–tail length (STL) of 5 cm. Juvenile axolotls measuring 5 cm STL and larger were fed commercial fish pellets daily until one year of age, after which feeding was reduced to three times per week. All animals were housed at a constant temperature of 18–20 °C.

### Limb and tail amputations

Juvenile animals measuring 5 cm snout–vent length (SVL) or sexually mature animals aged 2 years were used for all experiments. Prior to surgery, axolotls were anesthetized in a 0.01% benzocaine solution. Depth of anesthesia was verified by gently pinching a hindlimb to confirm the absence of a movement response. Amputations were performed on a plastic surface under an Olympus SZX10 stereomicroscope. For forelimb and tail amputations, the appendage was positioned flat on the surface, and a single perpendicular incision was made using a scalpel. Forelimbs were amputated at the midpoint of the humerus. Tail amputations were performed at 3 cm posterior to the cloaca in 5-cm SVL animals and at 9 cm posterior to the cloaca in 2-year-old animals. Following limb amputation, muscles were allowed to retract naturally, and the exposed bone was trimmed using scissors and forceps to create a flat amputation plane. After surgery, animals were returned to clean water, maintained at 20 °C, and allowed to recover with normal feeding.

### EdU pulse labeling

EdU was prepared as a 2.5 mg/mL stock in DMSO and administered via intraperitoneal injection at a dose of 2.5 µL per gram of body weight. Two hours upon EdU administration, animals were euthanized by anesthetic overdose using 0.1% benzocaine.

### Live imaging during regeneration and regenerate length measurements

Leucistic *d/d*, *Rag1*-KO, or *Foxn1*-KO axolotls and corresponding sibling controls were anesthetized in a 0.01% benzocaine solution prior to each imaging session. Anesthetized animals were positioned in Petri dishes coated with jellified 1% agar, positioning the regenerating limb or tail flat against the surface. Live imaging was performed using a Zeiss Axio Zoom V16 stereomicroscope. Regenerating limbs or tails were imaged approximately three times per week for juvenile animals, and once per week for sexually mature animals.

The size of regenerating limbs or tails in the different mutant contexts was measured from the amputation plane with Fiji. For statistical analyses, unpaired, two-tailed Student’s T-tests were performed between knock-out and the corresponding wild-type samples for each time point.

### Sample collection

#### Cryosectioning

Samples were collected in ice-cold PBS/4% PFA, fixed overnight at 4°C and washed in PBS overnight at 4°C. Samples were transferred to PBS/30% Sucrose and left to sink at 4°C overnight. Samples were embedded in OCT (Cryochrome^TM^ Blue) and kept at −20°C until sectioning. Sections of 10- or 12-µm-thickness were collected onto Superfrost-plus microscopy slides and air-dried for 1 hour before storing at −20°C until usage.

#### Whole-mount immunofluorescence or skeletal stainings

Samples were collected in ice-cold PBS/4% PFA, fixed overnight at 4°C and washed in PBS overnight at 4°C. Samples were stored at 4°C in clean PBS after fixation until further usage for maximum 1 week.

#### Spleen collection and weighing

Spleens were collected, rinsed with PBS, and weighed immediately after collection.

#### RNA extraction, cDNA synthesis and RT-qPCR

Samples were collected, rinsed with PBS, and transferred into ice-cold TRIzol, and kept at −80°C until RNA extraction. For RNA extraction, samples were homogenized with a hand-held KONTES tissue grinder. Phenol/chloroform RNA extraction was performed, and RNA was stored at −80°C until further use. For cDNA synthesis, RNA was treated with Ambion DNase I and 500 ng were used as template for cDNA synthesis using the iScript (Bio-Rad) cDNA Synthesis Kit according to the manufacturer’s instructions.

Primers for RT-qPCR were designed to amplify approximately 150 bp of the mRNA sequence of each target gene. RT-qPCR reactions were set up using the iQ SYBR Green Supermix (Bio-Rad) with 500nM forward and reverse primers each. All reactions were run for at least 3 biological replicates and in technical triplicates. The ΔCq values were normalized with the Cq value of EF1α. Negative controls (without cDNA template) were used in every RT-qPCR experiment, and Cq values of those controls were used as minimum gene expression detection threshold. For statistical analyses, unpaired, two-tailed Student’s T-tests were performed between the knock-out biological replicates and the corresponding wild-type controls of the corresponding experiment. Primers used for RT-qPCR are listed in Supplementary Table 2.

### Histology and stainings

#### *In situ* hybridization (ISH) on axolotl cryosections

The coding sequence for each target gene was mapped in the axolotl transcriptome version Am_3.4. Primers used to clone the ISH probes are listed in Supplementary Table 1. Probes were cloned in pGEMT-easy vectors and sequenced. Probes were transcribed with Digoxigenin (DIG)-labeling mix, purified through LiCl precipitation and diluted at 660ng/mL in hybridization buffer as working solution for ISH.

The ISH procedure was performed on cryosections as described before (Harland, 1991; Knapp et al., 2013). Briefly, cryosections were washed in PBS with 0.1% Tween20 and permeabilized with PBS/0.3% Triton-X100 for 20 minutes at room temperature (RT). Sections were hybridized with DIG-labeled probes in hybridization buffer (50% formamide, 10% dextran, 5X saline-sodium citrate solution (SSC), 0.1% Tween, 1mg/ml yeast tRNA, 1X

Denhardt’s solution, 0.1% CHAPS, 5mM EDTA in DEPC-treated de-ionized H2O) overnight at 60°C. Sections were then washed with wash buffer (50% formamide, 2X SSC, 0.1% Tween in DEPC-treated de-ionized H2O) twice for 30 minutes each at 60°C, washed with 0.2X SSC, 0.1% Tween for 30 minutes at 60°C and again for 30 minutes at RT. Sections were treated with 20µg/ml RNAse A in TNE buffer (10mM TRIS pH7.5, 500mM NaCl, 1mM EDTA in DEPC-treated de-ionized H2O) for 15 minutes at RT. The probe was detected with alkaline phosphatase (AP)-conjugated anti-DIG Fab fragments (Roche, 1:5000) through an antibody staining overnight at 4°C and BM purple (Roche) incubation at RT. The slides were mounted in PBS/50% Glycerol, sealed, and then imaged with a color camera using a Zeiss Axio Zoom V16 stereomicroscope.

#### Whole-mount RNA *in situ* hybridization of zebrafish fins

Whole-mount RNA in situ hybridization was carried out to assess the levels of *marco* in *rag1*^T26683^ zebrafish using a probe against *marco* (ENSDARG00000059294) (in courtesy of Alejandra Cristina López-Delgado). The oligonucleotides 5’-GGATATTTAGGTGACACTATAGAAAGGGGAAAGAGGGATGGTTG -3’ (forward) and 5’-GGATTAATACGACTCACTATAGGGCTGTGCATGGAGAACTGACA -3’ (reverse) were first used to amplify cDNA sequences to obtain an 860-base pair fragment. The PCR was run with DreamTaq reagents (Thermo Scientific) according to the manufacturer’s instructions and the products were purified using NucleoSpin Gel and PCR Clean-up Kit (Macherey-Nagel, Thermo Scientific). The *marco* probe was then transcribed *in vitro* using T7 RNA Polymerase (Roche, # 10881767001) according to manufacturer’s instructions. The reaction mixture was then treated with DNaseI (Invitrogen, # 18068015) according to the manufacturer’s instructions to eliminate residual DNA. The probe was then precipitated overnight at −80°C after the sequential addition of Lithium Chloride (100 mM) and absolute ethanol (3 times the reaction volume). The following day, supernatant was removed after centrifugation for 15 minutes at 4°C (13000 rpm) to obtain the pellet. It was then washed with 75% ethanol, followed by centrifugation (15 min at 4°C, 13000 rpm) and supernatant removal. The pellet was finally dried and resuspended in nuclease free water and stored at −20 °C after the addition of equal volume of formamide. It was to be diluted at 1:400 for further use.

On day 1, samples were gradually rehydrated using 75 %, 50 %, and 25 % methanol solutions in PBT (PBS with 0.1 % Tween) for 5 min each, followed by four washes in 100% PBT. Then, Proteinase K (Roche, REF:03115879001) was used (20 min in 20 μg/ml ProtK in PBT). The digest was stopped by rinsing 2 times in PBT, followed by refixation in 4% PFA in PBS (15 min) and 5 washes in PBT for 5 minutes each at room temperature. Samples were incubated in the hybridization buffer [5x SSC (saline-sodium citrate), 500μg/ml type VI Torula yeast RNA (Sigma R6625) or type V tRNA from wheat germ (Sigma R7876), 50μg/ml Heparin, 0.1% Tween 20, 9 mM citric acid (monohydrate) pH 6.0 - 6.5, 50% formamide (deionized)] for 1.5h at 65°C in a water bath and were left to hybridize overnight at 67°C with 200μl of the preheated probe diluted (1:400) in the hybridization buffer. On day 2, after recovering the probe, washing was done in a 67°C water bath, with all solutions pre-heated to the same temperature. The samples were washed in the hybridization buffer for 20 minutes, 3 times in 50% 2x SSCT [saline sodium citrate with 0.1 % Tween 20)/ 50% deionized formamide (PanReac AppliChem ITW Reagents, #A2156,1000)] for 20 minutes each, 2 times in 2x SSCT for 20 minutes each and 4 times in 0.2x SSCT for 30 minutes each. After washing the samples in PBT for 5 minutes at RT, they were incubated in blocking solution [5% sheep serum (Sigma S2263), 10mg/ml BSA (bovine serum albumin) in PBT)] for 1 hour at RT. Finally, samples were incubated overnight in anti-Dig-AP antiserum (Roche, REF:11093274910, 1:2000) in PBT + 2mg/ml BSA at 4°C. On day 3, the antibody was recovered, and samples were washed 2 times 5 min each, followed by 6 washes of 20 minutes each in PBT. Then, staining buffer NTMT was prepared freshly (100 mM Tris HCl pH 9.5, 50 mM MgCl2,100 mM NaCl, 0.1% Tween 20) and used to wash the samples 3 times (5 min each). The samples were then stained using 500μl NBT/BCIP staining solution (20μl NBT/BCIP in 1 ml NTMT - Roche, REF:11681451001) in the dark. Signals were monitored using a dissecting microscope, and the staining reaction was stopped by rinsing two times in PBT and adding STOP solution (0.05M phosphate buffer pH 5.8, 1mM EDTA, 0.1% Tween 20) upon achieving the desired signal strength. After rinsing 2 times in PBT, ethanol clearing was carried out, in which the samples were washed two times in 100 % ethanol for 15 min, once in 50 % ethanol for 5 min and 4 times in PBT for 5 min. Finally, they were kept in 80 % glycerol / 20 % STOP solution till imaging.

#### Embedding and cryosectioning of fins

Fins of *rag1*^T26683^ mutant fish were washed in PBS 2 times (20 minutes each) at RT and left in 0.5 M EDTA overnight at 4°C. The fins were then washed in 10 %, 20 % and 30 % sucrose solutions in PBS and 30 % sucrose in PBS and OCT (Tissue-Tek, REF:4583) mix (1:1) for 30 minutes each at RT. The fins were incubated overnight in OCT at 4°C and frozen into cryo-blocks after appropriate placement, followed by the labelling of the block. The embedded fins were stored in an ultra-freezer (−70 °C) until sectioning. The CryostarTM NX70 cryostat (Thermo Scientific) was used to section the embedded fins at 12μm thickness using Superfrost plus slides, which were stored at −20°C until use.

#### Whole-mount immunofluorescence coupled with tissue clearing

Whole-mount stainings were performed as previously described using the Salamander-Eci method (Subiran Adrados et al., 2021). Samples were incubated at room temperature (RT) on a rocker from the permeabilization step and until before dehydration. Samples were permeabilized in permeabilization solution (0.3M Glycine, 2% Triton X-100, 20% DMSO in PBS) for 3 overnights. Samples were blocked with blocking solution (1.5% inactivated goat serum, 10% DMSO, 2% Triton X-100 in PBS) for 1 hour. For EdU-incubated samples, Click-iT Plus EdU Cell Proliferation Kit (Invitrogen) was used to prepare the EdU staining solution according to the manufacturer’s instructions, however, the fluorescent azide was diluted 30 times from the recommended usage. Samples were incubated in the EdU staining solution for 5 hours. Samples were protected from light from this step onwards. Samples were washed in washing buffer (2% Tween-20, 1% heparin in PBS) overnight and blocked in blocking solution for 4 hours. Samples were incubated with primary antibodies diluted in washing buffer during 3 overnights and washed for one day in washing buffer. Samples were incubated with secondary antibodies diluted in washing buffer for 4 hours. For the nuclear staining, samples were incubated in SYTOX Green Nucleic Acid Stain (Thermofisher, 1:5000) diluted in 10X washing buffer for 1 hour and washed with washing buffer for one day. Samples were dehydrated through serial Ethanol dilutions in PBS at 4°C without rocking. Samples were incubated in 30%, 50% and 70% Ethanol solutions for 2 hours each, and in 100% Ethanol overnight at 4°C. Samples were cleared by incubating in Ethyl cinnamate (Sigma Aldrich) at RT and kept at RT until imaging. Antibodies used are listed in Supplementary Table 3.

Anti-CD2 or anti-CD79A stainings with EdU were analyzed with Arivis Vision 4D software V 4.1.1. A pipeline was generated for antibody- and/or EdU-positive cell quantification and volume determination of the regenerating structures. Threshold values of the pipeline were adjusted with 5 randomly chosen images from each antibody staining. For statistical analyses, unpaired, two-tailed Student’s T-tests were performed between each time point and the untouched limb.

Thymic nodule volumes were analyzed in Fiji with the plugin Volumest (http://lepo.it.da.ut.ee/~markkom/volumest/). For statistical analysis, unpaired, two-tailed Student’s T-tests were performed between the Rag1-KO and wild-type nodules.

#### Immunohistochemistry (IHC) on slides

Incubations were done at room temperature unless stated otherwise. Slides were air-dried for 15 minutes, washed with PBS for 3 minutes and incubated in PBS/3% H2O2 for 5 minutes. Slides were washed with PBS for 10 minutes, permeabilized with PBS/0.2% Triton X-100 for 15 minutes and then blocked for 1 hour with blocking buffer (PBS/0.3% Triton X-100/10% inactivated goat serum). Slides were then incubated with primary antibodies in blocking buffer overnight at 4 °C in a humidified chamber. Afterwards, slides were washed with PBS for 10 minutes and then with PBS/0.1% Triton X-100 for 5 minutes. Slides were incubated with secondary HRP-conjugated antibodies diluted in blocking buffer for 1 hour, rinsed with PBS and washed with PBS for 5 minutes. Slides were incubated for 10 minutes in freshly prepared peroxidase substrate (DAB Substrate Kit, Vector Labs) according to manufacturer’s instructions. Slides were rinsed and washed with de-ionized H2O for 5 minutes, washed with PBS for 10 minutes and mounted with PBS/50% Glycerol. Slides were imaged with a Zeiss Axio Zoom V16 stereomicroscope.

#### Immunofluorescence (IF) on slides

Incubations were performed at room temperature unless stated otherwise. Slides were air-dried for 15 minutes, permeabilized with PBS/0.2% Triton X-100 for 30 minutes and blocked for 1 hour with blocking buffer (same as buffer used for IHC). Slides were incubated with primary antibodies in blocking buffer overnight at 4 °C in a humidified chamber. Slides were washed with PBS/0.2% Triton X-100 for 30 minutes, incubated with secondary antibodies diluted in blocking buffer for 4 hours and washed in PBS/0.3% Tween-20 for 10 minutes. Slides were incubated with Hoechst 33342 (Invitrogen, 10 mg/mL stock in HPLC H2O, 1:10000) or SYTOX Green (Thermofisher, 1:5000) diluted in PBS/0.3% Tween-20 for 15 minutes, washed with PBS for 20 minutes and mounted with DAKO fluorescence mounting medium. Slides were imaged using a Zeiss Axio Zoom V16 fluorescence stereomicroscope.

#### Immunohistochemistry on tail fin sections

Immunofluorescence was carried out to assess the number of L-plastin positive cells and Alcama expression in the regenerate. For the staining steps, humidified staining chambers were used, while aluminium covered Coplin jars were used for washing steps. Slides were first washed three times for 5 min each in PBT (PBS with 0.1 % Tween-20), followed by incubation in blocking buffer [10% goat serum (Gibco) in PBT] for 1 hour at 4 °C. Parafilm was used to cover the slides for an even distribution (for other incubation steps as well). Meanwhile, the primary antibody solution was prepared by adding rabbit-anti-L-plastin (1:500, in courtesy of the Michael Brand lab, CRTD) and mouse-anti-Zn-8/Alcama/Neurolin (1:200, DSHB, # AB_531904) to blocking buffer. After removal of the blocking solution, the samples were incubated with the primary antibodies overnight at 4 °C. The following day, slides were washed in PBT for 6×10 minutes and incubated with secondary antibodies in blocking buffer for 45 minutes at room temperature. The secondary antibody solution was prepared by adding goat-anti-rabbit Alexa 488 (1:500, Thermo Fisher, # A11034) and goat-anti-mouse Alexa 555 (1:500, Thermo Fisher, # A21424), respectively, to blocking buffer.

Slides were then washed with PBT 3×10 min and stained with 4’,6-diamidino-2-phenylindole (DAPI) in PBT (1:40000 of 1mg/ml stock) for 5 minutes. After washing 3×10 minutes with PBS, the slides were mounted and sealed using Vectashield Hardset antifade mounting medium (Vector Laboratories). Slides were further stored at 4 °C for imaging and analysis.

#### Skeletal stainings

Fixed limbs in PBS were dehydrated through serial Ethanol dilutions in PBS at 4°C without rocking. Samples were incubated in 30%, 50% and 70% Ethanol solutions for 2 hours each, and in 100% Ethanol overnight at 4°C. Samples were transferred to pure ice-cold acetone and incubated for 2 overnights with gentle rocking. Samples were incubated in skeletal staining solution (0.03% alcian blue, 0.005% alizarin red, 5% acetic acid, 70% Ethanol in de-ionized H2O) for 1 week at RT with gentle rocking. Samples were washed in acetic acid solution (5% acetic acid, 70% Ethanol in de-ionized H2O) for 1 hour, and in 95% Ethanol at 4°C overnight. Samples were cleared by washing in 3% KOH solution (20% glycerol, 3% KOH in de-ionized H2O) for 1 week changing the solution every other day. Samples were transferred to 25% glycerol in PBS and imaged with a Zeiss Axio Zoom V16 stereomicroscope.

The length of the skeletal elements was measured manually with Fiji. For statistical analyses, unpaired, two-tailed Student’s T-tests were performed between knock-out and the wild-type samples for each condition.

#### Naphthol AS-D Chloroacetate (NCAE) staining for granulocytes

For detecting and analysing neutrophils in *rag1*^T26683^ mutant fish population, the Leukocyte Naphthol AS-D Chloroacetate (Specific Esterase) Kit (Sigma-Aldrich 91C-1KT) was used on 12 µm sections of homozygous, heterozygous and wild type *rag1* mutant fish (sections courtesy of Marharyta Hnatiuk).

Initially, all the components of the staining kit were warmed at room temperature for 20 minutes, while deionized water was prewarmed at 37°C in a water bath (maintained throughout the protocol). The slides were air-dried for 20 minutes, followed by rehydration using PBS for 10 minutes at RT, to remove the OCT medium. Meanwhile, NCAE staining solutions were prepared strictly following the order of addition (for 4 slides: 0.25mL sodium nitrite solution, 0.25mL Fast Red Violet LB base, 10mL pre-warmed water, 1.25mL TRIZMAL 6.3 buffer concentrate, 0.25mL Naphtol AS-D chloroacetate solution). Then, sodium nitrite solution was added to the respective base solution in a Falcon tube. It was mixed and allowed to stand for 2 minutes. Pre-warmed water, TRIZMAL buffer concentrate and the substrate solutions were added, turning the solution red (NCAE). The staining solution was then added to Lockmailer tubes containing slides and were incubated in the water bath at 37°C for 15 minutes covered from light. The slides were then rinsed in deionised water for 2 min at RT, followed by nuclear staining with 4’,6-diamidino-2-phenylindole (DAPI) in PBT (1:1000) for 5 minutes. After washing for 3×10 minutes with PBS, the slides were mounted and sealed using the Vectashield Hardset antifade mounting medium. Slides were stored at 4 °C for imaging and analysis.

### CRISPR/Cas9-mediated mutagenesis

#### gRNA design and synthesis

Rag1 was mapped in the axolotl genome v3.0.0 (BioNano) and orthology to human and mouse genes was assessed using BLAST against the NCBI database. Introns, exons, as well as the coding sequence were mapped using the transcriptome version Am_3.4. crRNAs were designed with the Broad Institute CRISPick tool (Doench et al., 2016) and aligned against the axolotl genome to ensure the uniqueness of the sequences. Four crRNA sequences were cloned into the DR274 plasmid (Hwang et al., 2013) and gRNAs were synthesized using the MEGAshortscript T7 Transcription Kit (Invitrogen) according to the manufacturer’s instructions. gRNAs were purified by LiCl precipitation, diluted to a 5 µg/ µL stock and stored at −80°C until usage. All the crRNA sequences used are listed in Supplementary Table 4. The injection mixture consisted of 5 µg guide RNA (gRNA) and 5 µg Cas9 protein (inhouse) in a final volume of 10 µL RNase-free water. The injection mixture was incubated at 37 °C for 5 min prior to microinjection.

#### Microinjections

Microinjection of one-cell–stage leucistic *d/d* axolotl embryos was performed as previously described (Khattak et al., 2009). Embryos were collected in fresh tap water, briefly rinsed in 70% ethanol, and washed thoroughly with tap water. Embryos were manually dejellied using forceps in 1× Marc’s Modified Ringer’s (MMR) solution supplemented with penicillin/streptomycin (1× MMR, 13% pen/strep) under an Olympus SZX10 stereomicroscope. Dejellied embryos were transferred to 1.5% agar-based injection molds prepared with injection solution (1× MMR, 28% pen/strep, 20% Ficoll). Embryos were microinjected and allowed to recover for 2 h in fresh injection solution before being transferred to 1× MMR, 13% pen/strep, 5% Ficoll and incubated overnight. All microinjection procedures were done at 20°C.

#### Axolotl genotyping

Injected embryos or embryos from mutant founders were allowed to develop at 18 to 20°C for 3 weeks, when tail clips were performed for genotyping and analysis of their mutations. Genomic DNA was prepared using the Mouse Direct PCR Kit (Biotool) according to manufacturer’s instructions. Genomic target regions were amplified by PCR with the 2X M-PCR OPTI Mix (Biotool) using primers with the following annealing sequences: Rag1_g386-Fwd: GTCTATCAAAACTGACAACAACCAG Rag1_g386-Rev: GTGTCCACATCACTTCTCACGTC Amplicons were barcoded for each animal sample and pooled for deep-sequencing through Illumina short-read, paired-end sequencing. The generated datasets were analyzed using the CRISPResso2 tool (Clement et al., 2019) to identify frame-shift mutations and the percentages of these mutations per animal. Due to mutagenesis efficiency, only mutants generated with the gRNA386 were used for the characterizations shown in this work. Further, only F1 or F2 animals were used for experiments.

#### Zebrafish genotyping

400 μl of freshly prepared lysis buffer (100 mM Tris HCl pH 8-8.5, 200 mM NaCl, 0.2 % SDS, 5 mM EDTA pH 8, deionised water and 100 μg/ml Proteinase K) was added to fin clips and samples incubated in a Thermoblock for 1 hour (800 rpm, 55°C), followed by centrifugation for 10 minutes (14800 rpm). The supernatant was transferred to a fresh tube, and an equal volume of isopropanol was added, mixed and centrifuged (10 min at 14800 rpm). The supernatant was discarded, and the pellet was washed with 70% ethanol, followed by a spin down (5 min at 14800 rpm). The supernatant was discarded, and the pellet was dried and dissolved in deionised water.

For breeding purposes and analyses, *rag1* zebrafish were genotyped. After isolating DNA from fin clips, the sequence of interest with the point mutation was PCR amplified using the *rag1*^t26683/t26683^ specific primers 5’-TTTGTGACTCAACACGTGCTG-3’ and 5’- CGACTCGGATTAGGCTTCTG -3’ (Wienholds et al., 2002). The PCR was run with DreamTaq reagents (Thermo Scientific) according to the manufacturer’s instructions and the products were purified using NucleoSpin Gel and PCR Clean-up Kit (Macherey-Nagel, Thermo Scientific). Then, 15 µl of purified PCR products, diluted to 1 ng/µl along with 2 µl of 10 μM forward or reverse primer primers, were sent for sequencing by Eurofins Genomics Germany GmbH.

#### Imaging of zebrafish fins, image processing and analysis

The fin regenerates of *rag1*^T26683^ mutant zebrafish were imaged with a Zeiss Discovery V12 stereomicroscope with a 1.5x objective at 50x magnification using the brightfield channel. The microscope was equipped with an AxioCam MRm and AxioVision software version 4.7.1.0. The fins after whole-mount in situ hybridization were imaged using the same microscope at 60x magnification. Images of the sections after immunostaining and NCAE staining were taken using a Zeiss AxioImager ApoTome2 upright motorized microscope in the widefield mode. The same magnification was used for all the images [20x (N.A. 0.8) and 40x (N.A. 0.95)]. Images were processed and/or analyzed using the ZEN software version 2.6. or using Affinity Photo and ImageJ/Fiji version 1.54f (Schindelin *et al*., 2012). Fin regenerate length was measured using the Fiji line tool on the outer fin rays 2 to 4. The same tool was used to measure the Alcama+ domain length after immunostaining on fin sections. The area and density of the Alcama+ domain was calculated with the help of the Fiji polygon selection tool. The Fiji cell counter plugin was used to count L-plastin+ cells in a 100 μm x 100 μm region on the sections. and to analyse the number of neutrophils per section after NCAE staining. Affinity Photo version 1.10.5.1342 was used to generate figures with identical brightness and contrast settings. Finally, unpaired t-tests (two-tailed) with Welch’s correction were performed on datasets to compare two populations using GraphPad Prism version 9.5.2 and Brown Forsythe and Welch ANOVA test was performed to compare the datasets with three or more populations. A p-value less than or equal to 0.05 was considered significant for all tests.

### Bulk RNA sequencing

#### Sample collection and sequencing

Rag1-KO 1-year-old animals and leucistic *d/d* age-matched controls (both intact) were amputated on the same day and allowed to regenerate for 12 days. Forelimb mid-bud blastemas were collected in A-PBS (0.8X PBS). Three blastemas from different animals per condition were used and processed separately. Stump tissue was removed from the blastema with the help of forceps and scissors under an Olympus SZX10 stereomicroscope. Blastemas were washed gently in A-PBS and transferred to TRIzol for phenol/chloroform RNA extraction.

Sequencing of the purified RNA was performed using Smart-seq2. Sequence alignment and differentially expressed genes (DEGs) analyses were done using the axolotl genome assembly AmexG_v6.0-DD for sequence alignment and annotation. Sequence and gene annotation of the axolotl nuclear genome assembly AmexG_v6.0-DD were downloaded from https://www.axolotl-omics.org. The gene models for the HoxA and HoxD genes were replaced with manually curated gene models (personal communication with Sergej Nowoshilow, axolotl-omics.org). Sequence and gene annotation of the mitochondrial genome assembly (NCBI GenBank, AY659991.1) and RNA spike-in control sequences (ERCC,

Ambion; for quality control) were included as well. Adapters and poly(A)-tails were trimmed from the raw reads using cutadapt (v4.2) (Martin et al., 2011) with settings “--minimum-length=35 --pair-filter=any -a nextera_i5=CTGTCTCTTATA -a polyA=AAAAAAAAAA -a polyT=TTTTTTTTTT -A nextera_i5=CTGTCTCTTATA -A polyA=AAAAAAAAAA -A polyT=TTTTTTTTTT --times=2”. Trimmed RNA-seq reads were mapped against the reference using STAR (v2.7.10b) (Dobin et al., 2013) and splice sites information from the gene models. Uniquely mapped reads were converted into counts per gene and sample using featureCounts (v2.0.1) (Liao et al., 2014) and settings “-s 0 -p -Q 1”.

##### DEG analysis

Normalisation of raw fragments based on library size and testing for differential expression between the different conditions was done with the DESeq2 R package (v1.38.2, R v4.2.0) (Love et al., 2014). Principal Component Analysis based on the 500 most variable genes was performed to explore the relationship between biological replicates and different libraries. Differentially expressed genes were determined using the Wald test of DESeq2. Resulting p-values were corrected for multiple testing with the Independent Hypothesis Weighting package (IHW 1.26.0) (Ignatiadis et al., 2016). Genes with a maximum of 5% false discovery rate (adjusted p-value ≤ 0.05) were considered significantly differentially expressed.

##### Mapping and Counts

The scRNA-seq data was converted into counts per gene and cell using Cell Ranger (v7.1.0)[https://www.10xgenomics.com/]. To build the required Cell Ranger index, reference and annotation was first compiled as described in the methods section for the analysis of the bulk RNA-seq data. Since some tools of the Cell Ranger pipeline cannot deal with chromosomes longer than ∼536.9Mb due to their implementation, reference sequences and annotation were split into 500Mb chunks. The index was then build using the ‘mkref’ command of Cell Ranger.

#### Gene ontology (GO) enrichment analysis

Significantly up- or downregulated differentially expressed genes (DEGs) were used for GO enrichment analysis. All axolotl DEGs annotated with identifiers different as those used for human genes were aligned against the NCBI Gene database to find potential human orthologs. Analysis was done using the human database in the GOnet website https://tools.dice-database.org/GOnet/ (Pomaznoy et al., 2018). Only 281 of the 630 downregulated, and 42 out of the 182 upregulated DEGs could be mapped to the human database for the analysis. Heatmaps were plotted using the https://www.bioinformatics.com.cn/en online platform.

## List of supplementary materials

Supplementary Figures. S1 to S6

Supplementary Tables 1 to 4

## Acknowledgments

We thank Beate Gruhl and Anja Wagner (TU Dresden) for axolotl husbandry, Dunja Knapp for help with CRISPR-KO design, and all members of our individual laboratories for critical input. We are grateful to the Tübingen 2000 Screen Consortium (*Max-Planck-Institut für Entwicklungsbiologie, Tübingen:* Busch-Nentwich, E., Dahm, R., Frank, O., Frohnhöfer, H.-G., Geiger, H., Gilmour, D., Holley, S., Hooge, J., Jülich, D., Knaut, H., Maderspacher, F., Maischein, H.-M., Neumann, C., Nicolson, T., Nüsslein-Volhard, C., Roehl, H., Schönberger, U., Seiler, C., Söllner, C., Sonawane, M., van Bebber, F., Wehner, A., Weiler, C.) and Exelixis Deutschland GmbH (Erker, P., Habeck, H., Hagner, U., Hennen, Kaps, C., E., Kirchner, A., Koblizek, T., Langheinrich, U., Loeschke, C., Metzger, C., Nordin, R., Odenthal, J., Pezzuti, M., Schlombs, K., deSantana-Stamm, J., Trowe, T., Vacun, G., Walderich, B., Walker, A., Weiler, C.) for sharing *rag1* mutant zebrafish. We thank Michael Schorpp for providing zebrafish, and M. Fischer and S. Kunadt for excellent fish care. This work was supported by the Histology, LMF, Dresden Concept Genome Centre and FACS core facilities of the CMCB Technology Platform at the TU Dresden.

## Author contributions

L.B.C., Y.O.V., F.K. and M.H.Y. designed experiments. L.A.B.C., Y.O.V. performed experiments. A.P. pre-processed RNA-seq data. L.A.B.C., Y.O.V., F.K. and M.H.Y. interpreted and discussed results. M.H.Y. wrote the manuscript with input from all authors.

M.H.Y. and F.K. supervised the project.

---

**Supplementary figure 1.**
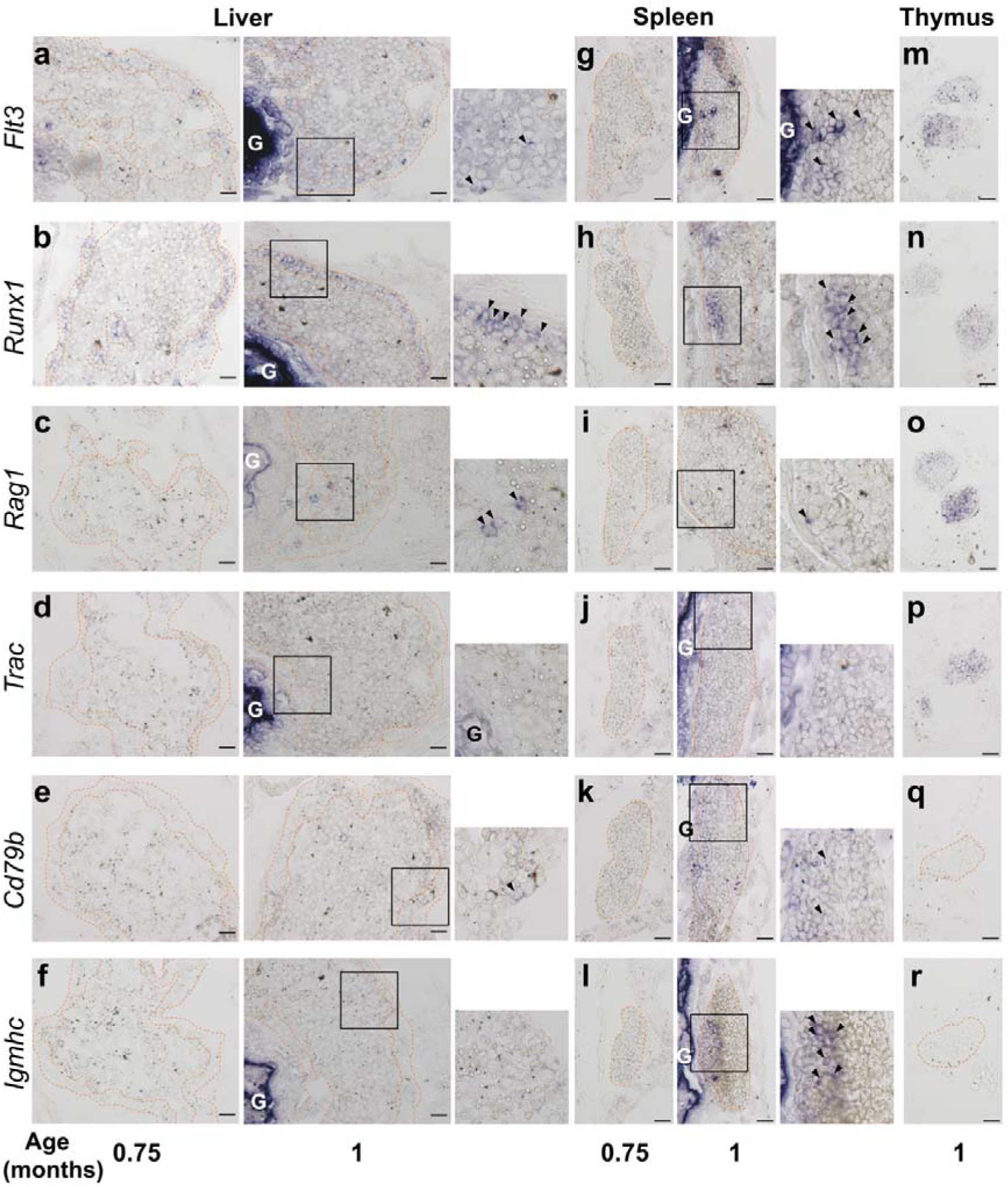
Markers of hematopoiesis and lymphopoiesis are expressed in axolotl tissues within one month of age. a-r) *In situ hybridization* (ISH) on cryosections from whole 3- or 4-weeks-old larvae. Liver cortical area, spleen and thymus in panels (q) and (r) are delineated with an orange dashed line. Positive cells are indicated with arrowheads at insets from 4-weeks-old larvae. G: gut. N=3, scale bar = 50 μm.

**Supplementary figure 2.**
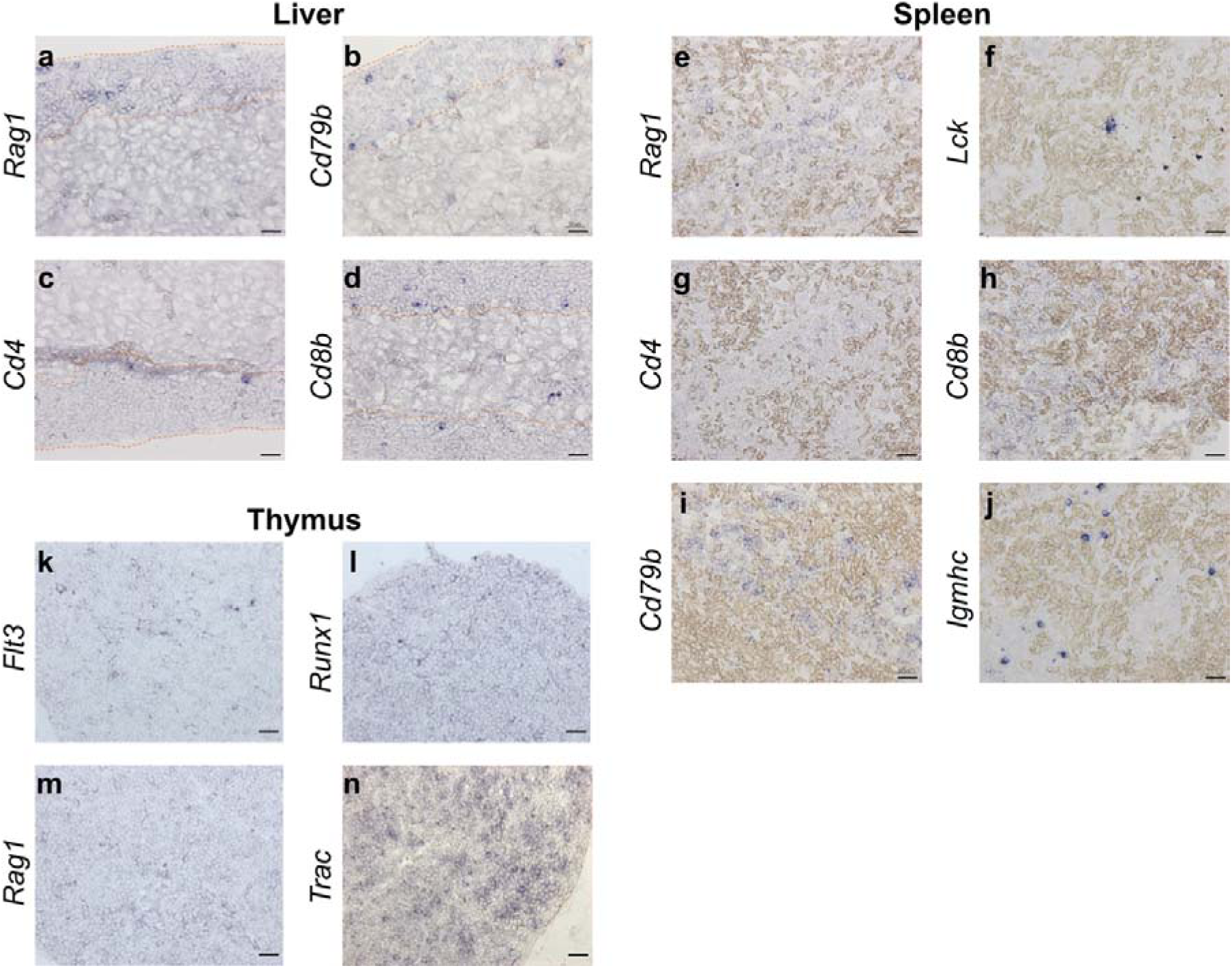
Markers of hematopoiesis and lymphopoiesis are expressed in tissues from sexually mature axolotls. a-j) ISH on cryosections from tissues of 1-year-old axolotls. The liver cortical area is delineated with orange dashed lines. N=3, scale bar = 50 μm.

**Supplementary figure 3.**
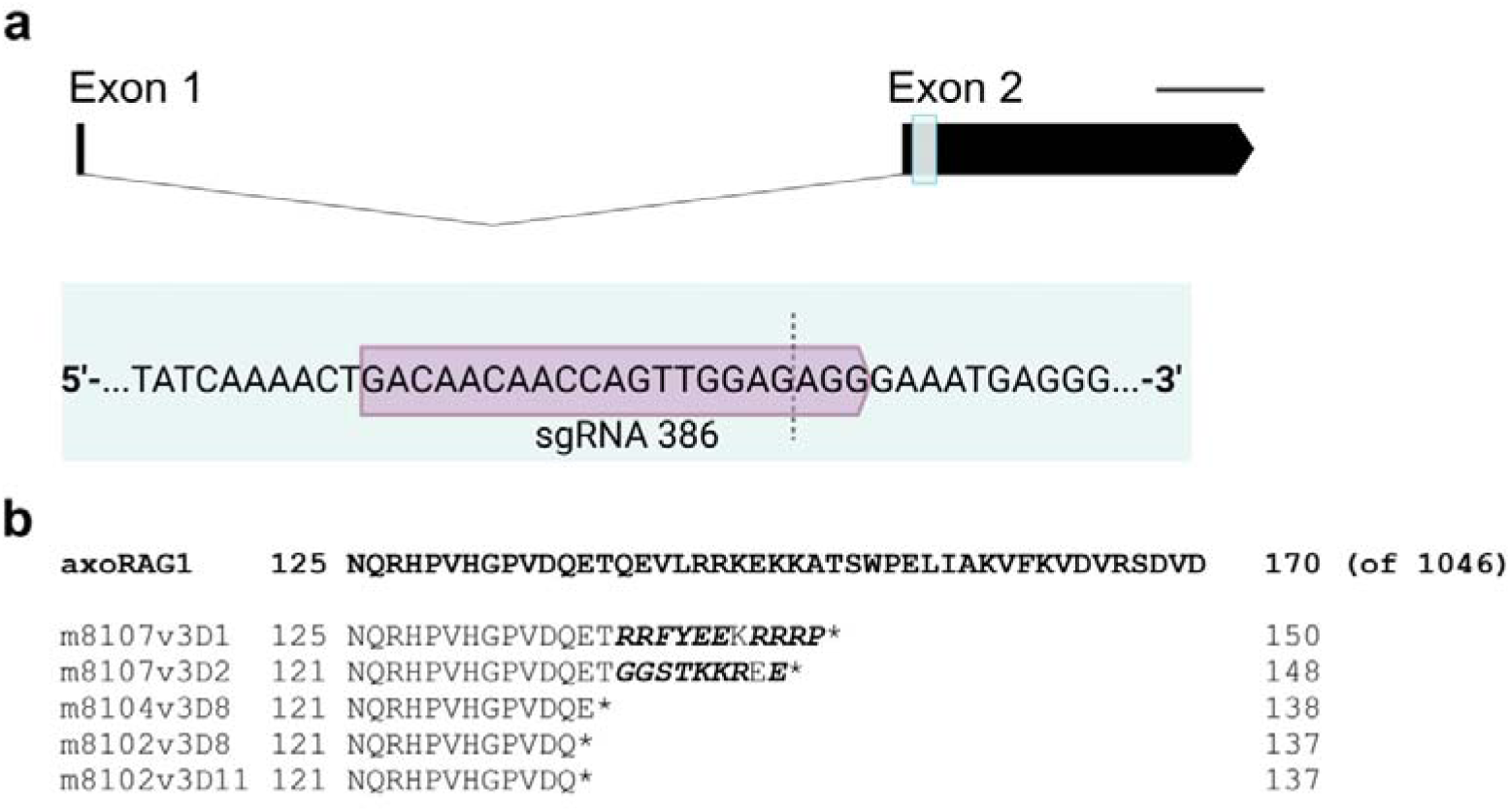
Generation of a Rag1 knockout axolotl line. a) *Rag1* gene structure, comprised by 2 exons, highlighting the region targeted by gRNA386. Expected cutting site is indicated with a dashed line. Scale bar = 1kb. b) RAG1 amino acid sequences obtained via CRISPR/Cas9-mediated mutagenesis. The position of the mutation within the gene is indicated (ranging from position 8102 to 8107), together with the version of the genome used for the annotation (v3) and the number of deleted nucleotides for each case (ranging from 1 to 11). Amino acid sequences are aligned against the wild-type sequence. Changes in amino acids are indicated in bold italics and stop codons with an asterisk. The position of the early stop codon for each generated mutant is indicated in the right column.

**Supplementary figure 4.**
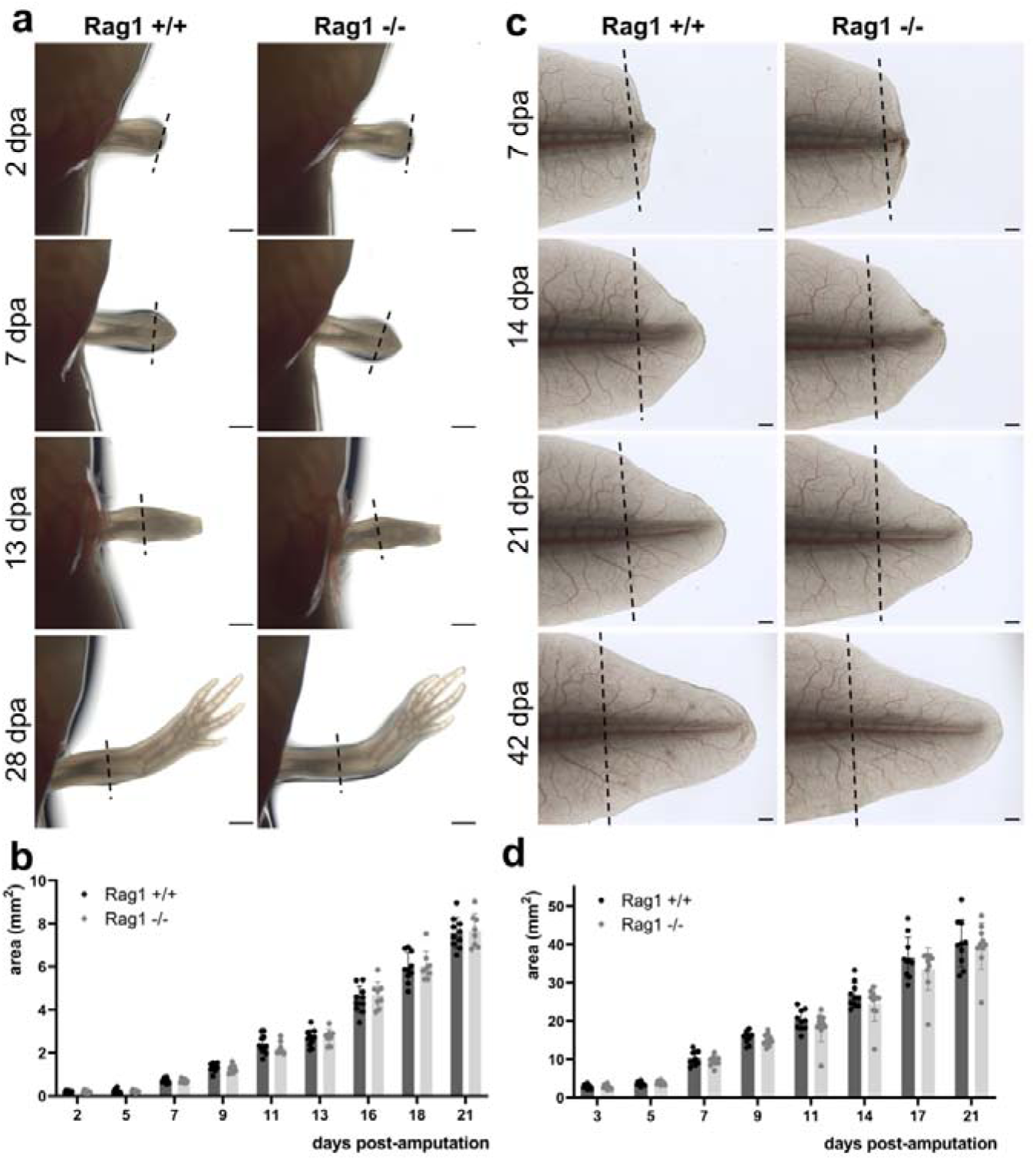
T and B lymphocyte depletion does not delay appendage regeneration in juvenile axolotls. a) Limb regeneration in juvenile wildtype and Rag1 −/− axolotls. Scale bar = 1 mm. b) Area of regenerated limb tissue (n=10, N=2, mean with SD). c) Tail regeneration in juvenile wildtype and Rag1 −/− axolotls. Scale bar = 1 mm. d) Area of regenerated tail tissue (n=10, N=2, mean with SD). Unpaired Student’s T-test performed for statistical analyses against the wildtype data in (b,d). No statistical significance observed.

**Supplementary figure 5.**
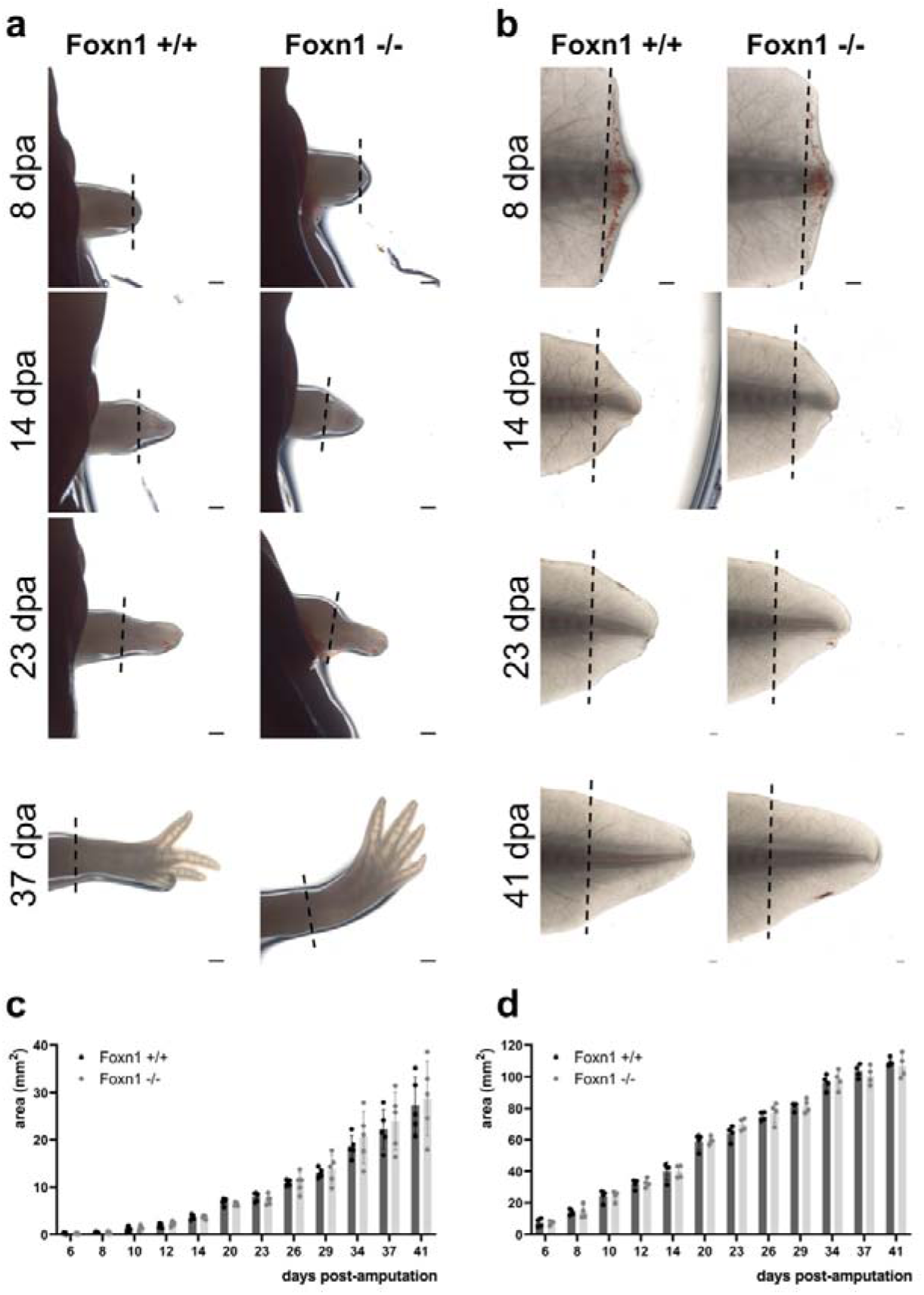
T lymphocyte-depletion does not delay appendage regeneration in axolotl. a,b) Limb (a) and tail (b) regeneration in juvenile wildtype and Foxn1 −/− axolotls. Scale bar = 1 mm. c,d) Area of regenerated limb (c) or tail (d) tissue (n=5, N=2, mean with SD). Unpaired Student’s T-test performed for statistical analyses against the wildtype data in (c,d). No statistical significance observed.

**Supplementary figure 6.**
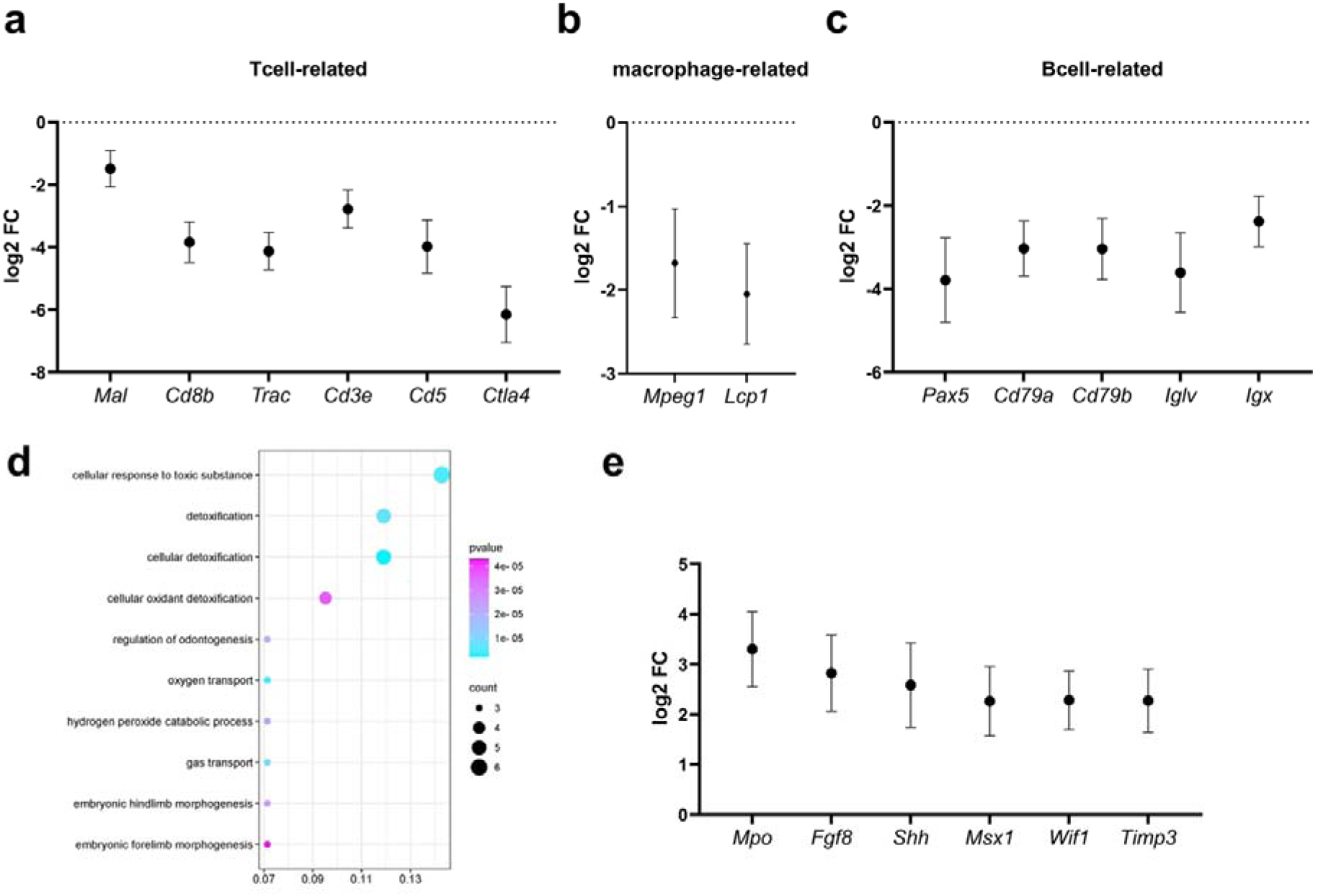
Lymphocyte depletion yields transcriptional changes in regenerating limbs. a-c) Significantly downregulated genes related to specific immune cell-type functions. d) GO enrichment analysis on upregulated DEGs. e) Significantly upregulated genes, related to neutrophil populations, limb morphogenesis and ECM remodeling. Plots represent the average log2 of fold change and bars show standard error (n=3). Only DEGs with an adjusted p value < 0.05 are shown.

